# *Aeschynomene americana* induces terminal bacteroid differentiation in *Bradyrhizobium* sp. USDA3516, a novel model for Dalbergioid-rhizobia symbiosis

**DOI:** 10.1101/2025.11.05.682141

**Authors:** T. Scott Carlew, Annika A. Atherton, Ashley Shim, Camilo Parada Rojas, Riley A. Buchanan, Jeff H. Chang, Joel L. Sachs, Brittany J. Belin

## Abstract

The paradigms of legume-rhizobia symbiosis are derived primarily from conserved features of Inverted-Repeat Lacking Clade (IRLC) legumes and closely related species. The Dalbergioids diverged from the IRLC early in legume evolution and possess unique symbiotic features but few genetically tractable models. The small, diploid Dalbergioid *Aeschynomene americana* (American jointvetch) has promise as a genetic model for Dalbergioid-rhizobia symbiosis, yet only a few studies have examined its symbiotic properties.

We examined the symbiont range of *A. americana* from central Florida and characterized a native *A. americana* nodule isolate, *Bradyrhizobium* sp. USDA3516.

We find that *A. americana* forms effective symbioses with *B.* sp. USDA3516, which is closely related to Thai *A. americana* symbiont *B.* sp DOA9, and with symbionts from the Dalbergioids stylo and peanut. Interestingly, several strains that effectively nodulated *A. americana* exhibited branched bacteroid morphologies, but we found that branching was neither necessary nor sufficient for effective symbiosis.

Our study contradicts the prevailing view that bacteroid shape is a major determinant of symbiotic efficiency and presents the *A. americana*-*B.* sp. USDA3516 interaction as an optimal model of *A. americana* symbiosis.

## INTRODUCTION

The Fabaceae or legume family of plants contains nearly 30,000 accepted species (Legume Phylogeny Working Group (LPWG), 2025), and many of these species form mutualistic symbioses with nitrogen-fixing rhizobia. Studies of the legume-rhizobia symbiosis have focused on the Papilionoideae subfamily, which is commonly split into Inverted-<u>R</u>epeat <u>L</u>acking <u>C</u>lade (IRLC) tribes and non-IRLC tribes based on presence of an inverted repeat (IR) region in their plastid genomes (Wojciechowski *et al*., 2004). Legume genetic tools first were developed in IRLC and closely related non-IRLC species, and as a result, the paradigms for legume-rhizobia symbiotic mechanisms are derived from conserved features across these clades.

According to these paradigms, legume-rhizobia symbiosis is initiated by Nod factors that induce development of infection threads and new meristem tissue in the root cortex. This meristem differentiates into nodule cells that are intracellularly infected by rhizobia, which in turn differentiate into bacteroids through a host-dependent process. In soybean, bacteroid differentiation is minimal and consists of upregulation of nitrogen fixation and metabolite exchange factors (Pessi *et al*., 2007). In *Medicago* and other genera, terminal bacteroid differentiation occurs that includes altered bacteroid morphology and increased endoreduplication. This terminal bacteroid differentiation is driven by host <u>n</u>odule-specific, <u>c</u>ysteine-rich (NCR) peptides (Guerra-Garcia & Sankari, 2025).

Dalbergioid legumes such as peanut, the world’s second most important commercial legume by production volume (Nations, 2024), diverged from the IRLC ancestor early in Papilionoideae evolution (**Fig. 1A**). This tribe is known to have unique symbiotic features. Many Dalbergioid legumes do not require Nod factors to initiate symbiosis (Giraud *et al*., 2007; Guha *et al*., 2022), and bacterial infection occurs through cracks in the root epidermis rather than an infection thread (Bonaldi *et al*., 2011; Noisangiam *et al*., 2012; Guha *et al*., 2022). Terminal differentiation of bacteroids does occur in Dalbergioids and is driven by host-produced peptides, but these peptides generally are longer and more anionic than IRLC NCR peptides (Czernic *et al*., 2015; Gully *et al*., 2018; Raul *et al*., 2022; Boukherissa *et al*., 2025). Identifying the molecular mechanisms of these unique symbiotic features of the Dalbergioids is key to understanding legume-rhizobia symbiosis evolution.

**Figure 1.**
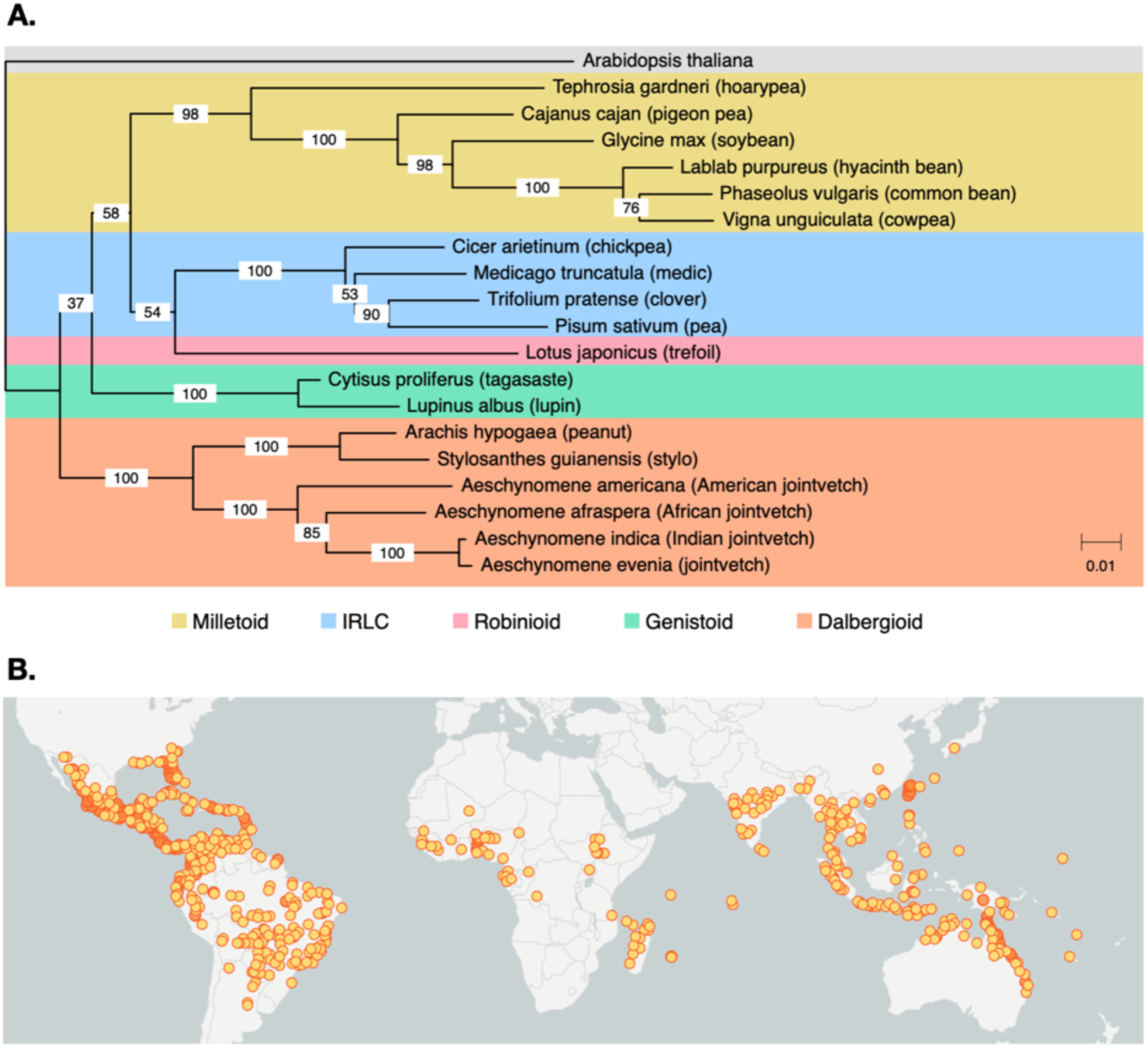
**Selected tribes of the *Papillionideae***. (**A**) Phylogenetic tree based on conserved regions of *matK* sequences. Scale bar indicates substitutions per 1000 nucleotides. (**B**) Geographic distribution of *Aeschynomene americana* (Legume Data Portal, Accessed Sept. 2025).

Most genetic work in the Dalbergioids has been performed in the jointvetches (*Aeschynomene* spp.), a genus of semi-aquatic legumes that includes the species *A. afraspera*, *A. indica*, *A. evenia*, and *A. americana*. Of these, only the diploid species *A. evenia* has a fully assembled genome (Quilbé *et al*., 2021). *A. americana* is also diploid and thus a promising alternative model for Dalbergioid genetics (Chaintreuil *et al*., 2016; Brottier *et al*., 2018), and it has commercial relevance as a cover crop and for rice paddy intercropping (**Fig. 1B**). Currently there are few studies on *A. americana* symbiosis. Prior work indicates that *A. americana* symbioses are infection thread-independent and that compatible symbionts (namely *Bradyrhizobium* sp. DOA9) encode genes for Nod factor production, are non-photosynthetic, and do not terminally differentiate (Noisangiam *et al*., 2012; Teamtisong *et al*., 2014; Okazaki *et al*., 2015).

Here, we examine the symbiotic compatibility of *A. americana* with a panel of rhizobium type strains, including the *A. americana* nodule isolate *B.* sp. USDA3516 (Grant & Trese, 1996). We affirm that *A. americana* is not compatible with photosynthetic *Bradyrhizobium* spp. and that its overall symbiont range is more like the Dalbergioids peanut and stylo. Contradicting earlier work, we do observe hallmarks of terminal bacteroid differentiation, including endoreduplication and altered bacteroid morphology; however, altered morphology was not observed in all productive strains. Sequencing of *B.* sp. USDA3516, the most effective strain tested, indicates that this strain likely produces Nod factors and encodes the BclA protein involved in NCR import (Guefrachi *et al*., 2015). These findings suggest that production of NCRs to induce terminal bacteroid differentiation may be universal in the Dalbergioids and is thus an ancient symbiotic mechanism.

## RESULTS

### *A. americana* symbiont range

Previous work indicated that *A. americana* nodulation is Nod factor-dependent and that *A. americana* is not nodulated with photosynthetic *Bradyrhizobium* spp. (Noisangiam *et al*., 2012). To investigate its symbiont range more broadly, we inoculated *A. americana* from central Florida with a collection of 13 rhizobium type strains, as well as one native strain (*B.* sp. 3516) that isolated from root nodules of *A. americana* grown in the same region (full strain names are provided in **Table 1**). The type strains used included strains engaged in both Nod factor-dependent and Nod factor-independent symbiosis from diverse hosts.

**Table 1.** Strains of rhizobia used for *A. americana* inoculation. . ‘^T^’ indicates type strains. ‘***’ indicates strain was natively isolated from *A. americana*.

| Symbiont Strain | Host Species | Host Common Name | Host Tribe |
| --- | --- | --- | --- |
| *** *Bradyrhizobium* sp USDA3516 | *Aeschynomene americana* | American jointvetch | Dalbergioid |
| *Bradyrhizobium* sp ORS285 | *Aeschynomene afraaspera* | African jointvetch | Dalbergioid |
| *Bradyrhizobium* BTAi1<sup>T</sup> | *Aeschynomene indica* | Indian jointvetch | Dalbergioid |
| *Bradyrhizobium arachidis* LMG26975<sup>T</sup> | *Arachis hypogaea* | Peanut | Dalbergioid |
| *Bradyrhizobium stylosanthis* BR10<sup>T</sup> | *Stylosanthes guianensis* | Stylo | Dalbergioid |
| *Bradyrhizobium diazoefficiens* USDA110<sup>T</sup> | *Glycine max* | Soybean | Milletoid |
| *Bradyrhizobium japonicum* USDA6<sup>T</sup> | *Glycine max* | Soybean | Milletoid |
| *Bradyrhizobium elkanii* USDA76<sup>T</sup> | *Glycine max* | Soybean | Milletoid |
| *Sinorhizobium fredii* NGR234<sup>T</sup> | *Glycine max* | Soybean | Milletoid |
| *Bradyrhizobium cajan* AMBPC1010<sup>T</sup> | *Cajanus cajan* | Pigeon pea | Milletoid |
| *Sinorhizobium fredii* HH103<sup>T</sup> | *Lablab purpureus* | Hyacinth bean | Milletoid |
| *Bradyrhizobium semiaridum* LMG31654<sup>T</sup> | *Tephrosia gardneri* | Hoarypea | Milletoid |
| *Bradyrhizobium brasilense* R<sup>T</sup> | *Vigna unguiculata* | Cowpea | Milletoid |
| *Bradyrhizobium canariense* BTA-1<sup>T</sup> | *Cytisus proliferus* | Tagasaste | Genistoid |

At 14 days post-inoculation, we quantified shoot height, nodule counts, nodule dry biomass, and acetylene reduction rates of plants inoculated with each strain (**Fig. 2**). Four strains provided robust benefits to *A. americana*, eliciting >10 nodules per plant on average, and had significantly higher nitrogenase activity than uninoculated control plants, including *B. stylosanthis*, *B. arachidis*, *B.* sp. USDA3516, and *B. cajani*. One strain, *Sinorhizobium fredii* HH103, provided significant fixed nitrogen but formed relatively few nodules. *B. diazoefficiens* inoculation increased plant shoot heights but produced few, ineffective nodules, suggesting that the benefit is not strictly due to symbiotic nitrogen fixation.

**Figure 2.**
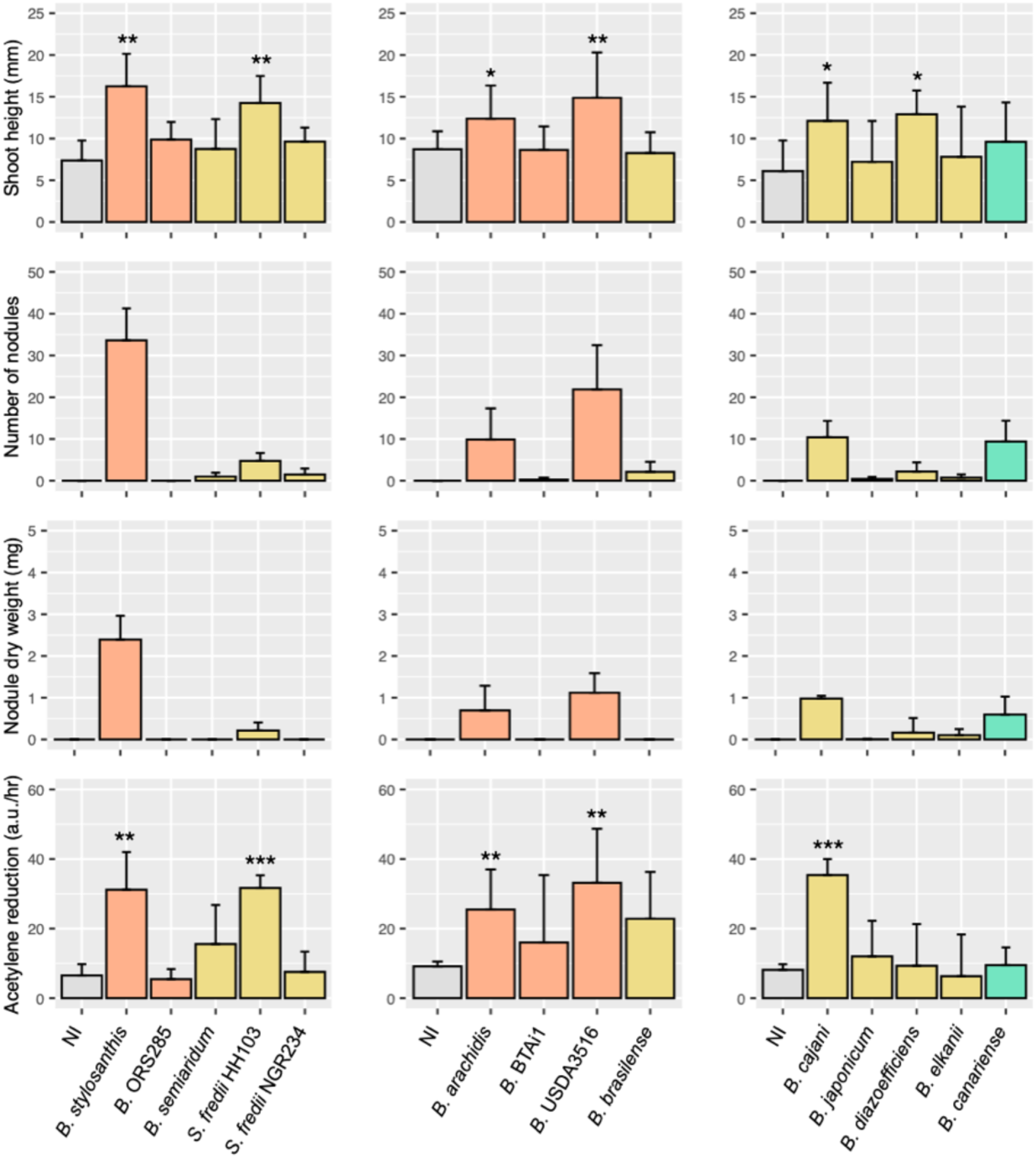
*A. americana* growth is promoted by non-photosynthetic Dalbergioid symbionts. Shoot height, nodule number, and nodule dry weight, and acetylene reduction activity for plants inoculated with each strain from Table 1. NI = ‘Non-Inoculated’ control. N=7-9 plants per condition. Error bars indicate standard deviation. Colors of bars indicate native host tribe, according to the legend in Figure 1. Asterisks indicate p-values from two-tailed t-test comparison with non-inoculated plants: ‘*’ < 0.01, ‘**’ < 0.001, ‘***’, < 0.0001.

None of the remaining strains conferred benefits to *A. americana*. *B. canariense* nodulated *A. americana* robustly, but these nodules did not fix significant nitrogen. *B. semiaridum*, *S. fredii* NGR234, and *B. brasilense* nodulated weakly but also were ineffective. None of the remaining strains formed nodules on *A. americana,* including *B.* sp. ORS285, *B.* BTAi1, *B. japonicum*, and *B. elkanii*.

### *A. americana* nodule and bacteroid morphology

We next examined nodule morphology from *A. americana* inoculated with each strain that exhibited robust nodulation (at least 10 nodules/plant on average), including *B. stylosanthis*, *B. arachidis*, *B.* sp. USDA3516, *B. cajani*, and *B. canariense*. We collected semi-thin sections of nodules harvested at 14 days post inoculation (dpi), stained with Calcofluor and the BacLight LIVE/DEAD kit, and imaged by confocal microscopy. Interestingly, bacteroids of all strains except for *B. cajani* appeared to have morphological changes relative to typical free-living *Bradyrhizobium* spp. (**Fig. 3**). Nodules containing *B.* sp. USDA3516 had much larger bacteroid volumes and branched bacteroid morphologies, whereas more subtle bacteroid morphological changes occurred in nodules with *B. stylosanthis*, *B. arachidis*, and the inefficient fixer *B. canariense*. We initially hypothesized that the variation in presence of morphological changes in bacteroids across strains was due to some strains requiring longer time windows to complete differentiation. However, nodules harvested at 35 dpi had similar bacteroid morphologies to the 14-dpi time point (**Fig. S1**).

**Figure 3.**
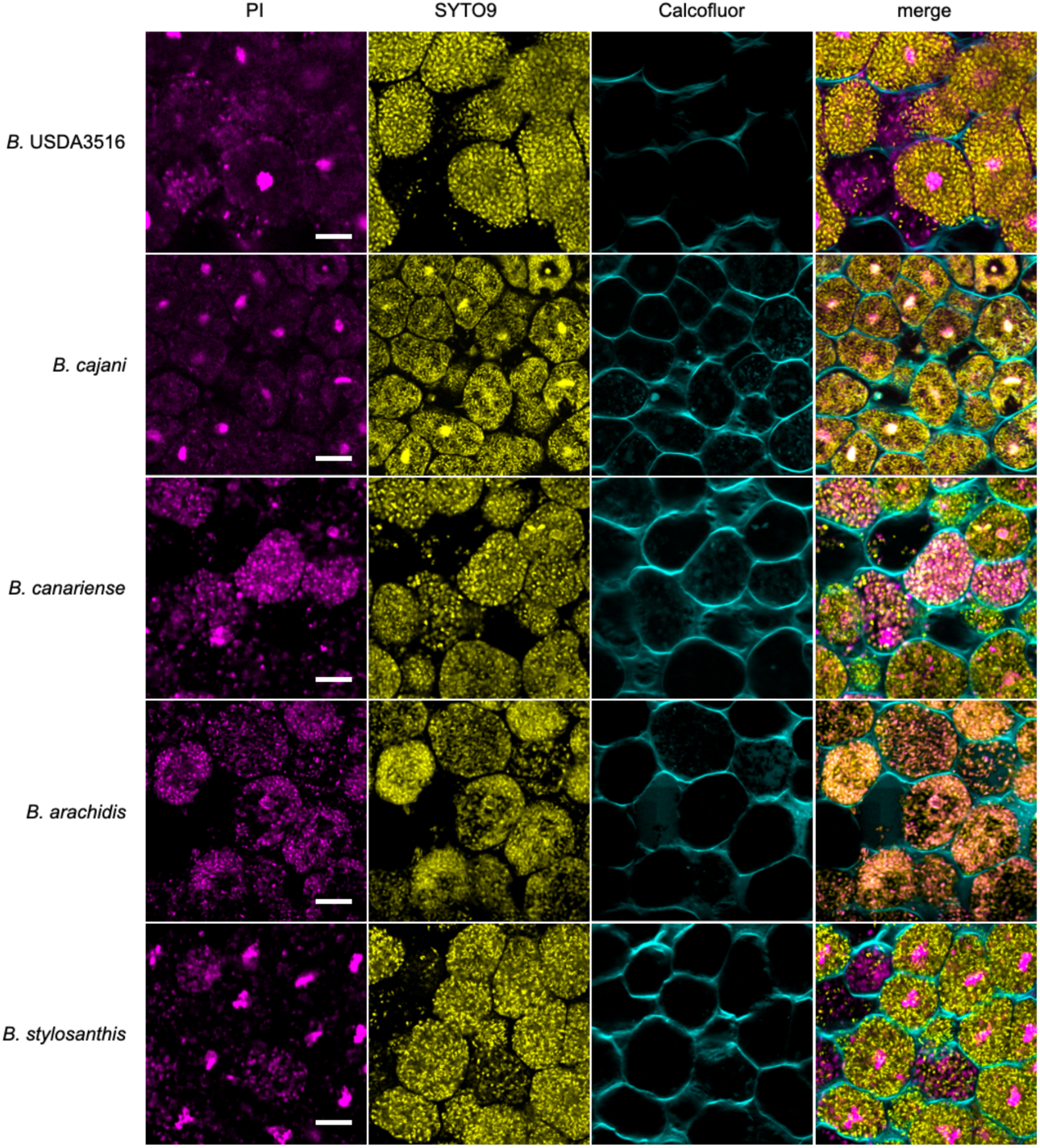
*A. americana* bacteroids are terminally differentiated. Representative confocal images of nodule cross- sections collected at 14 dpi. Sections were stained with Calcofluor (cyan), SYTO9 (yellow), and propidium iodide (“PI”; magenta). Scale bars = 10 microns.

To obtain more quantitative information on bacteroid characteristics, we isolated bacteroids from root nodules and compared them to free-living cells by imaging and flow cytometry (**Fig. 4**; **Fig. S2-S6**). In all strains, free-living cell morphologies were similar, exhibiting rod shapes of 3-6 micron length and 1-2 SYTO9 (DNA) foci in the cytoplasmic interior. Staining with the Nile Red (NR) dye under PHB granule-detecting conditions demonstrated that the free-living cells usually had few or no PHB granules. Bacteroid morphologies, however, were diverse across strains. In native symbiont *B.* sp. USDA3516 (**Fig. 4A**; **Fig. S2**), extracted bacteroids had a branched morphology and were slightly wider than free-living cells. Most bacteroids had multiple PHB granules and SYTO9 was either strong throughout the cytoplasm or present in multiple bright, discrete foci at the poles. The increased volume of *B.* sp. USDA3516 bacteroids was clear by forward scatter flow cytometry, and SYTO9 flow cytometry indicated that bacteroids had a higher ploidy than free-living cells, with a ∼20% increase in median SYTO9 fluorescence. Bacteroids of *B. stylosanthis* were similar in properties to *B.* sp. USDA3516, albeit with a slightly higher increase (40%) in median SYTO9 signal (**Fig. 4D**; **Fig. S6**). We verified that the apparent branched morphologies and larger areas of *B. stylosanthis* and *B.* sp. USDA3516 bacteroids were not due to clumping of multiple cells with the membrane dye FM 4-64, which revealed that branched bacteroids were surrounded by one continuous cell envelope (**Fig. S7-8**).

**Figure 4.**
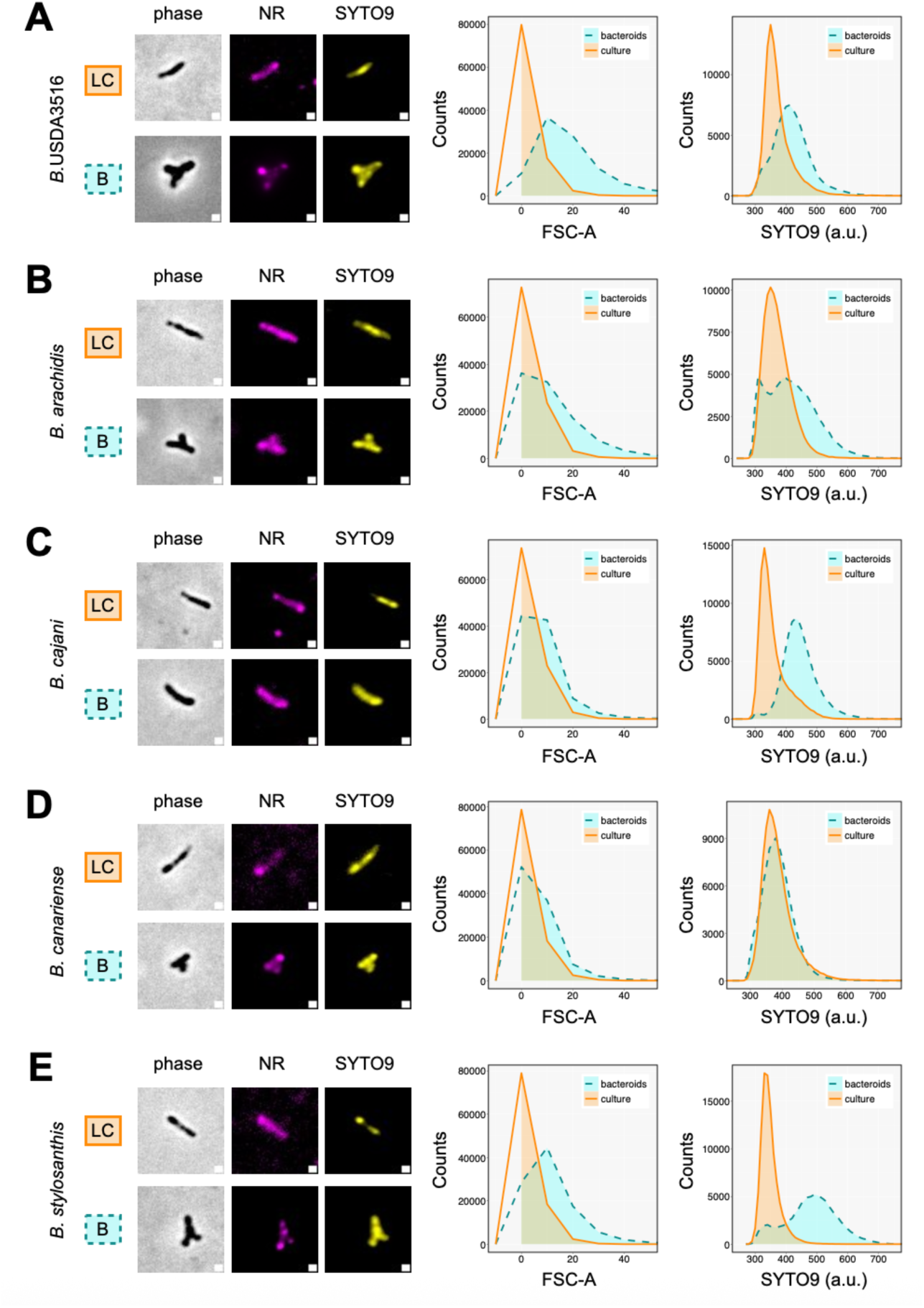
Productive bacteroids exhibit higher levels of endoreduplication. Representative phase and fluorescence images (*left*) with flow cytometry forward scatter distributions (*center)* and SYTO9 intensity distributions (*right*) for bacteroids (“B”, dotted blue filled lines) versus cultured cells (“LC”, solid orange filled line) after staining with Nile Red (NR) and SYTO9: (**A**) *B.* sp. USDA3516, (**B**) *B. arachidis*, (**C**) *B. cajani*, (**D**) *B. canariense*, (**E**) *B. stylosanthis*. Scale bars indicate 1 micron.

Differentiation of other strains was more variable. *B. arachidis* also formed mostly branched bacteroids with increased SYTO9 signals relative to free-living cells but contained less pronounced PHB granule staining than in *B. stylosanthis* and *B.* sp. USDA3516 (**Fig. 4B**; **Fig. S3**). Bacteroids of *B. cajani* were slightly larger and wider than free-living cells but were unbranched, and SYTO9 intensity relative to free-living cells was higher than in *B.* sp. USDA3516 (**Fig. 4C**; **Fig. S4**). Interestingly, bacteroids of *B. canariense* – the only ineffective symbiont to robustly form nodules – generally were branched and very slightly larger than free-living cells, but these bacteroids had no increase in ploidy. This suggests that endoreduplication rather than branching is the most important aspect of the differentiation, at least within our strain set.

### *Bradyrhizobium* sp. USDA 3516 genome properties

The apparent terminal differentiation of *B.* sp. USDA3516 in *A. americana* was surprising, given that terminal differentiation was not observed in the effective Thai *A. americana* isolate *B.* sp. DOA9 (Teamtisong *et al*., 2014). To investigate whether this difference could be explained by the gene content of the two strains, we sequenced and assembled the *B.* sp. USDA3516 genome. We found that the *B.* sp. USDA3516 genome contains a single 7.8 Mbp chromosome with 7,283 ORFs predicted by both Prokka and Bakta prokaryotic annotation tools (**Fig. 5**). We did not detect any native plasmids in this strain; commonly *Bradyrhizobium* spp. do not encode symbiotic plasmids (Weisberg, A *et al*., 2022).

**Figure 5.**
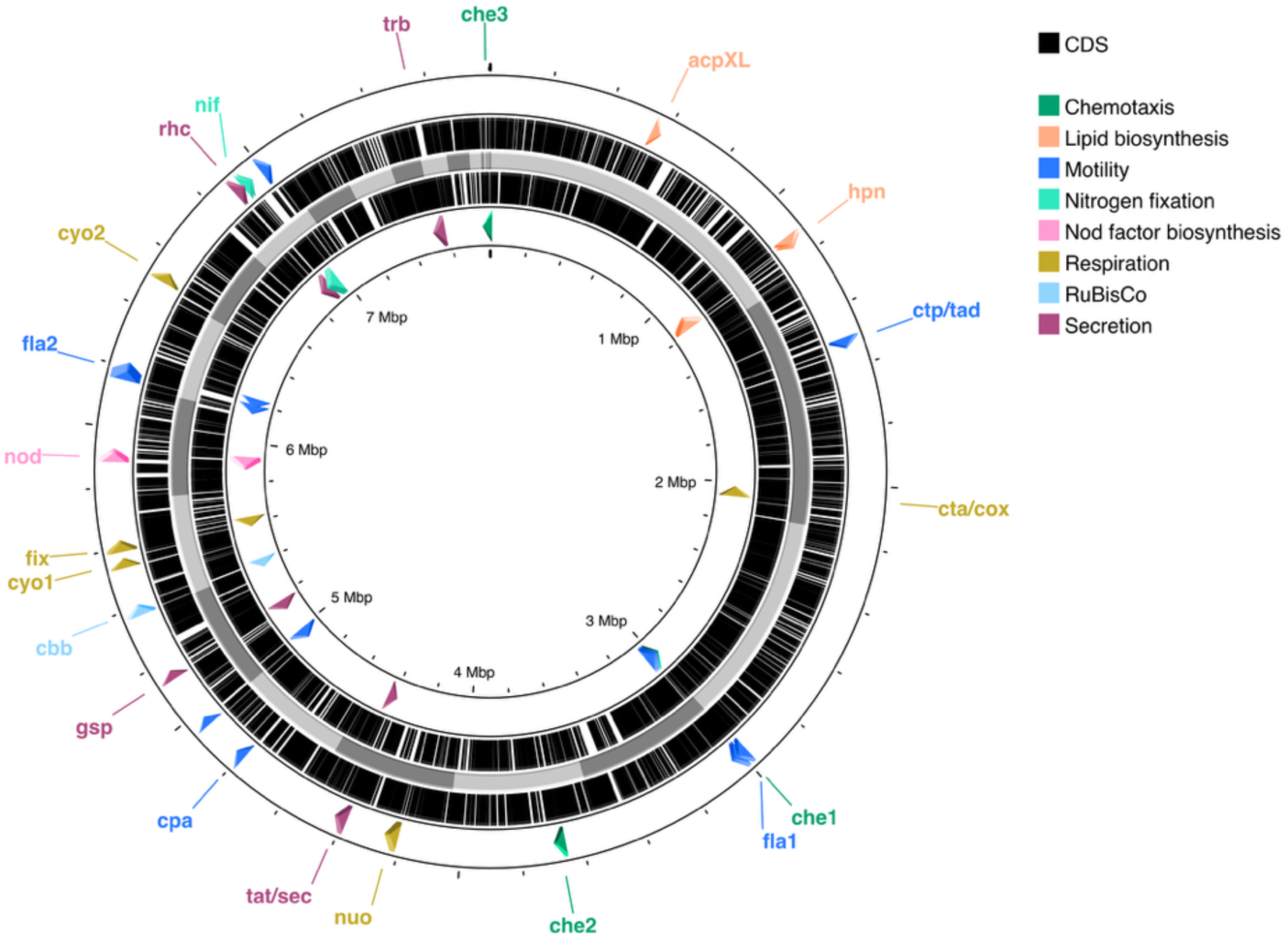
Genome architecture of native *A. americana* symbiont *B.* sp. USDA3516. Genome architecture of *B.* sp. USDA3516 consisting of a single circular chromosome. Inner ring (alternating grey regions) denotes individual scaffolds; second rings from center indicate positions of protein-coding genes (CDS, in black) as annotated in Table S1. Outermost rings from center indicate ORFs for symbiotically relevant gene categories, as denoted in the legend. Illustration generated using Proksee (https://proksee.ca/)(Grant et *al.*, 2023).

We annotated ORFs using the consensus from Bakta (Schwengers *et al*., 2021) and Prokka (Seemann, 2014) annotation tools, InterPro protein domain analysis (for protein-coding ORFs) (Jones *et al*., 2014), and reciprocal best BLAST hit analysis against closely related and model bacterial strains (list in Methods; **Table S1**). These revealed the presence of well-known symbiosis-driving genes, including *nod* genes, indicating that this strain can produce Nod factors (**Fig. 5**). The strain appears to be metabolically versatile with multiple respiratory terminal electron acceptors (*fix*/cbb3, *cox*/c1, and two *cyo/*bo3 operons), and though it contains a RuBisCo (*cbb*) gene cluster, it does not contain other genes for photosynthesis in bradyrhizobia (Mornico *et al*., 2012). It includes at least three secretion systems (*gsp*/T2SS, *rhc*/T3SS, *trb*/T4SS) and pili biosynthesis genes (*ctp*/*tad*), as well as two large flagellar gene clusters (*fla*) and three chemotaxis operons (*che*). Based on synteny with other Rhizobiales, we predict that flagellar cluster *fla2* encodes lateral flagellar genes (Garrido-Sanz *et al*., 2019). This organism also includes signature lipid species of *Bradyrhizobium* spp. and rhizobia, such as genes involved in the biosynthesis of hopanoid lipids (*hpn*) and the synthesis and addition of very long chain fatty acids to lipid A (*acpXL*).

### Comparison of *B.* sp. USDA 3516 to other symbiotic *Bradyrhizobium* spp

We next assembled a genome-wide phylogeny of *Bradyrhizobium* species representatives using bac120 alignments from the GTDB Toolkit (**Fig. 6**) (Chaumeil *et al*., 2022). As expected from the gene content, *B.* sp. USDA3516 is not part of the clade containing photosynthetic bradyrhizobia or the *B. japonicum* clade, and instead it is more closely related to other productive *A. americana* symbionts from this study: *B. cajani*, *B. arachidis*, and *B. stylosanthis*. The strain’s closest relatives are *B. yuanmingense* CCBAU 10071 and the Thai *A. americana* symbiont *B.* sp. DOA9, with which it shares >92% average nucleotide identity (**Fig. S9A**).

**Figure 6.**
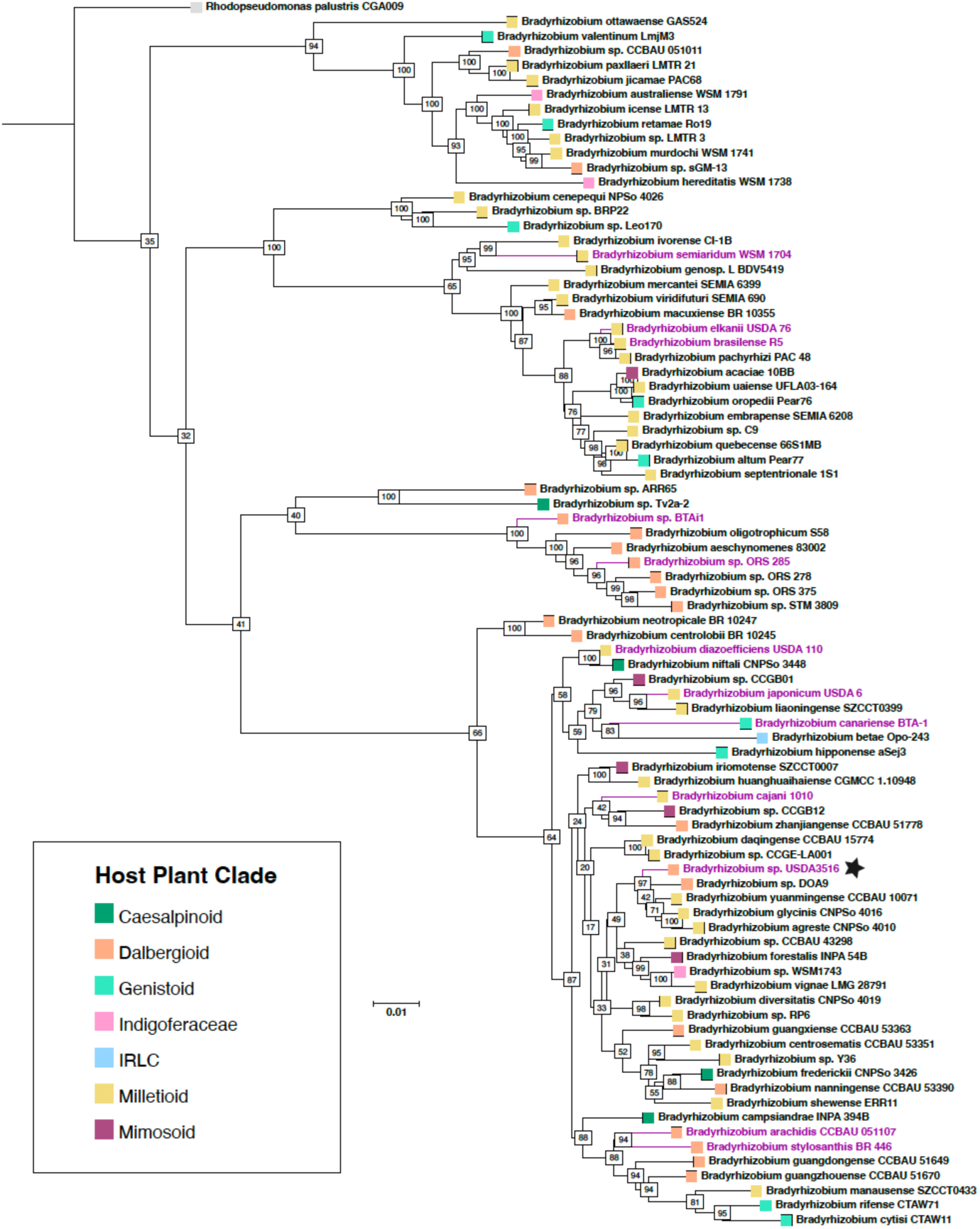
Bac120 species tree of representative *Bradyrhizobium* spp. Maximum likelihood tree for conserved regions across select *Bradyrhizobium* species representative. Branch names for species used for *A. americana* inoculation in this study are highlighted in purple, and the *B.* sp. USDA3516 genome is indicated with a black star. Colored boxes at branch tips indicate the legume clade (tribe/subfamily) of the host species from which the strain was originally isolated (see Legend). Numbers at internal nodes (highlighted in white) indicate branch support values using ultrafast bootstrap approximation (UFBoot), based on 1000 bootstraps: 100 = highest confidence, 0 = no confidence. The outgroup sequence is indicated with a grey branch tip label. Scale bar indicates substitutions per 1000 nucleotides.

The degree of nucleotide conservation between *B.* sp. DOA9 and *B.* sp. USDA3516 is surprising given their contrasting symbiotic phenotypes regarding bacteroid differentiation. We initially hypothesized that they may have lower than average similarity within genes known to be involved in the differentiation process. However, protein phylogenetic trees of the NCR peptide importer BclA (the *Bradyrhizbium* spp. homolog of BacA in *S. meliloti*) (Guefrachi *et al*., 2015) indicated that *B.* sp. USDA3516 BclA is more similar to the *B.* sp. DOA9 BclA than to the BclA proteins from other species with branched *A. americana* bacteroids (**Fig. S9B**). We also analyzed the FtsZ protein in *B.* sp USDA3516, based on recent data demonstrating that FtsZ depletion in the Rhizobiales/Hyphomicrobiales is sufficient to induce branching (Aubry *et al*., 2025), yet the FtsZ protein tree topology is likewise similar to the species relationships (**Fig. S9C**).

One distinction between *B.* sp. DOA9 and *B.* sp. USDA3516 that is not captured by comparing conserved genomic loci is the presence of unique gene groups, and it is possible that bacteroid branching in *B.* sp. USDA3516 is driven by genes that are not present in *B.* sp. DOA9. We used the Proteinortho tool (Klemm *et al*., 2023) to identify orthologous gene groups (OGs) found in all species with branched bacteroids (*B.* sp. USDA3516, *B. arachidis*, *B. stylosanthis*, and *B. canariense*) and in none of the non-branching species (*B.* sp. DOA9 and *B. cajani*) (**Fig. S9D**). This analysis yielded a list of 45 OGs (**Table 2**), which includes genes involved in LPS biosynthesis and DNA replication. Interestingly, seven of these OGs are part of a conserved gene cluster in that is adjacent to the MinCDE cell topology factors (**Fig. 7**). Though the Min system is not essential in rhizobia, misregulation of Min protein expression in *S. meliloti* can induce branching (Cheng *et al*., 2007), potentially indicating that altered regulation of the MinCDE region is required for branched bacteroids.

**Figure 7.**
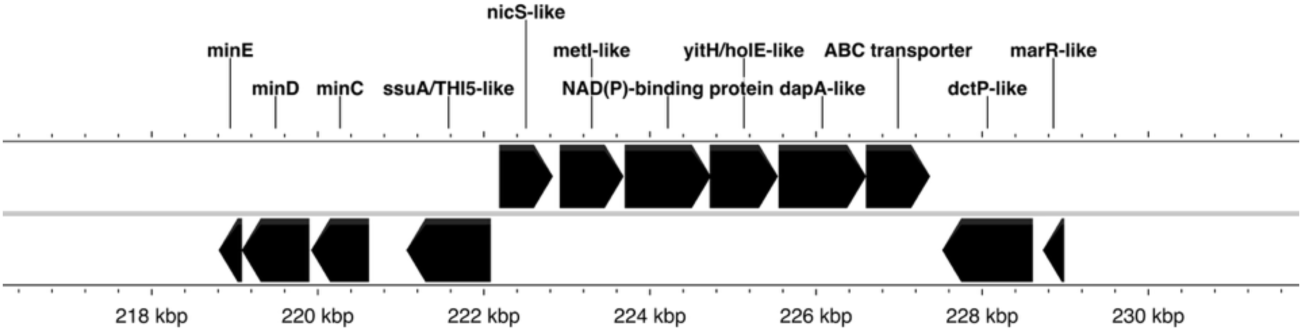
Conserved operon adjacent to *minCDE* in species with branched *A. americana* bacteroids. Illustration generated using Proksee (https://proksee.ca/)(Grant e*t al.*, 2023).

**Table 2.** Orthologous gene groups (OGs) specific to species with branched bacteroids in *A. americana*. OGs identified in all species with branched bacteroids (*B.* sp. USDA3516, *B. arachidis*, *B. stylosanthis*, and *B. canariense*) and in none of the non-branching species (*B.* sp. DOA9 and *B. cajani*). OGs in bold indicate a conserved operon.

| B. sp. USDA3516 locus | Gene | Description |
| --- | --- | --- |
| Brady3516_00244 |  | SsuA/THI5-like domain-containing protein |
| Brady3516_00245 |  | TetR family transcriptional regulator, NicS type |
| Brady3516_00246 |  | ABC transporter type 1, MetI-like superfamily |
| Brady3516_00247 |  | NAD(P)-binding domain superfamily |
| Brady3516_00248 |  | GNAT family N-acetyltransferase, YitH/HolE-like |
| Brady3516_00249 |  | DapA-like protein |
| Brady3516_00250 |  | Aliphatic sulfonates import ATP-binding protein SsuB-like |
| Brady3516_00266 |  | Ribonuclease Z/Hydroxyacylglutathione hydrolase-like metallo-beta-lactamase |
| Brady3516_00760 |  | Hypothetical protein |
| Brady3516_01459 |  | DUF6636 domain containing protein |
| Brady3516_01628 |  | AMP-dependent synthetase/ligase |
| Brady3516_02047 |  | NAD(P)-dependent dehydrogenase, short-chain alcohol dehydrogenase family |
| Brady3516_02208 | *fsr* | Fosmidomycin resistance protein |
| Brady3516_02463 |  | Hypothetical protein |
| Brady3516_02636 |  | AMP-dependent synthetase/ligase |
| Brady3516_03010 |  | Hypothetical protein |
| Brady3516_03026 |  | ABC-type multidrug transport system, ATPase component |
| Brady3516_03502 |  | Hypothetical protein |
| Brady3516_03543 |  | Hypothetical protein |
| Brady3516_03544 |  | CHRD domain-containing protein |
| Brady3516_03546 |  | Cyclic di-GMP phosphodiesterase |
| Brady3516_03565 |  | Hypothetical protein |
| Brady3516_03575 |  | Acetyltransferase, LpxA superfamily |
| Brady3516_03623 |  | Multidrug transporter EmrE superfamily, Amino Acid and Vitamin Transporters |
| Brady3516_03703 |  | Transmembrane protein TauE-like |
| Brady3516_03873 |  | Bordetella uptake gene |
| Brady3516_04043 |  | Epoxide hydrolase-like |
| Brady3516_04171 |  | ABC-type molybdate transport system, periplasmic Mo-binding protein ModA |
| Brady3516_04290 |  | Hypothetical protein |
| Brady3516_04336 | *pcaC_3* | 4-carboxymuconolactone decarboxylase/NDH-1 regulator |
| Brady3516_05191 |  | Hypothetical protein |
| Brady3516_05329 |  | DUF1236 domain-containing protein |
| Brady3516_06440 |  | Hypothetical protein |
| Brady3516_06649 | *rfbC* | dTDP-4-dehydrorhamnose 3,5-epimerase |
| Brady3516_06662 |  | Acetyltransferase, LpxA superfamily |
| Brady3516_06663 |  | DegT/DnrJ/EryC1/StrS family aminotransferase |
| Brady3516_06686 |  | Hypothetical protein |
| Brady3516_06716 |  | L-2-Haloacid dehalogenase |
| Brady3516_06994 |  | YjbJ superfamily |
| Brady3516_07077 | *traF* | Type IV secretory pathway, protease TraF |
| Brady3516_07079 | *repA* | Replication initiator protein A |
| Brady3516_07080 |  | DNA-binding protein |
| Brady3516_07083 |  | YspA, cpYpsA-related SLOG domain protein |
| Brady3516_07084 |  | TOPRIM domain protein |

## MATERIALS & METHODS

### Bacterial strain cultivation

Strain *Bradyrhizobium* sp. USDA3516 was obtained from the USDA ARS National Rhizobium Germplasm Collection, courtesy of Patrick Elia. *Bradyrhizobium* spp. ORS285 and BTAi1 were originally obtained from Eric Giraud (LSTM, Montpellier, France) and later gifted by Dianne Newman (Caltech). All other strains were purchased from the DSMZ-German Collection of Microorganisms and Cell Cultures.

All *Bradyrhizobium* spp. and *Sinorhizobium* spp. strains were streaked on rich AG medium agar plates (4.6 mM sodium gluconate, 6.6 mM arabinose, 1 g/L yeast extract, 6 mM NH4Cl, 5.6 mM MES, 5 mM HEPES, 1 mM Na2HPO4, 1.76 mM Na2SO4, 88 µM CaCl2, 25 µM FeCl3, and 0.73 mM MgSO4, pH 6.6) from 10% glycerol stocks. Plates were grown at 30℃ for 4-5 days before colonies were inoculated into 10mL AG rich liquid media. These cultures were grown to exponential phase (OD_600_ = 0.4-0.8) at 30℃ and 250 rpm prior to plant inoculation. For plant inoculation, cultures were pelleted by centrifugation at 4000g in a spinning bucket centrifuge and resuspended to OD_600_ = 1.0 in fresh AG. During inoculation, 300 uL of OD_600_ = 1.0 resuspensions were added to each 30 mL *A. americana* culture tube.

### A. americana cultivation

*Aeschynomene americana* seeds (Hancock Farm & Seed Company) were surface sterilized by a 5-minute incubation in 95% ethanol, followed by 30-minute incubation with 10% bleach, followed by 5 washes in sterile water. The seeds were then plated on 1% water agar plates. Plates were sealed with parafilm, wrapped to prevent light exposure, and germinated at 30℃ for 3 days. Germinated seedlings were rooted into 30 mL culture tubes filled with Bradyrhizobium Nodulation Medium (BNM). Rooted *A. americana* were then moved to growth chambers held at 12h light/dark cycles at 30℃ and 80% humidity for the duration of the experiment.

### Acetylene reduction assays

Whole *A. americana* plants were moved into 30 mL Balch tubes with 1mL of sterile water. Balch tubes were plugged with rubber septa and sealed with aluminum crimp caps, after which 10% of the headspace was removed via syringe and 16G needle and replaced with acetylene gas (Airgas). Samples were incubated overnight under growing conditions before measurement of ethylene production by GC-MS as described (Pan *et al*., 2024).

### Bacteroid extraction

Nodules were removed from roots and submerged in a 10% bleach solution for 5 minutes. Nodules were then washed three times with Bacteroid Extraction Buffer (BEB) (113.7mM Disodium Malate, 125 mM KCl, 50mM TES buffer) before homogenization with mortar and pestle in 5 mL of BEB. Samples were then centrifuged at 100g for 10 minutes to remove plant debris. Cleared supernatant was moved to a new tube, centrifuged at 1500 g for 20 minutes, and resuspended in 1 mL of ice-cold BEB. All subsequent steps were performed at 4°C with ice-cold buffers. The resuspension was gently added to the top of a freshly prepared Percoll gradient with 10 mL 85%/10 mL 60%/10 mL 45% Percoll layers. The sample was spun through the Percoll gradients 10,000g for 30 minutes, after which a band of concentrated bacteroids was visible. A 5 mL fraction containing this band was collected, diluted 1:10 in BEB, and then centrifuged at 1500g for 20 min. The bacteroid pellet was washed again in 5 mL BEB to remove residual Percoll, resuspended in PBS with 4% fresh paraformaldehyde, and fixed at 4°C overnight.

### Fluorescence staining

Nodule semi-thin sections (100 µm thickness) were collected using a 7000 smz-2 vibratome (Campden Instruments). Nodule sections were stained and mounted for imaging as described (Pan *et al*., 2024). For cultured samples, cells were grown to mid-exponential phase (OD_600_ = 0.6) and pelleted at 4000 x g for 30 minutes. Cell pellets were resuspended in PBS with 4% fresh paraformaldehyde and fixed at 4°C overnight. Fixed cultured cells and fixed bacteroids prepared as above were stained for microscopy by incubating with 7.5 μM SYTO9 for 30 minutes then with 0.25 μg/mL FM 4-64 for 10 minutes or with 0.5 μg/mL Nile Red for 30 minutes, all in PBS.

### Microscopy

Confocal fluorescence images of nodule cross-sections were taken using a Zeiss LSM980 confocal microscope using either a 40X/1.3 NA or 63X/1.4 NA objective. All images were collected using an Airyscan 2 detector with the following wavelength ranges for each dye: Calcofluor White, 405nm laser excitation and 422-477 nm emission; SYTO9, 488 nm excitation and 495-550 nm emission; Propidium Iodide, 561nm excitation and 607-735 nm emission. For imaging bacteroids, SYTO9 and FM 4-64 fluorescence images were acquired using a Zeiss LSM980 confocal microscope with a 40X/1.3 NA objective. Images were collected using an Airyscan 2 detector with the following wavelength ranges for each dye: SYTO9, 488 nm excitation and 483-506 nm emission; FM 4-64, 514 nm excitation and 500-751 nm emission. Phase images with SYTO9 and Nile Red fluorescence were acquired using a Nikon Ti2 inverted epifluorescence microscope with a 40X/0.75 NA phase objective. The following wavelength ranges were used for each dye: SYTO9 – 488 nm excitation, GFP emission filter cube (502-538 nm); Nile Red – 561 nm excitation, Nikon TRITC emission cube (570-613 nm).

### Flow cytometry

Cultured cells and bacteroids were prepared as described above, stained with 0.025 μM SYTO9 for 15 minutes in PBS, and fixed in 4% paraformaldehyde/PBS at 4°C overnight. Samples were washed three times in PBS and then diluted to 1:10 - 1:100 in 1 mL PBS, depending upon the sample density. Flow cytometry was performed on samples with an Attune Nxt Acoustic Focusing Cytometer rujning Attune Nxt Software v. 3.2.1526.0. Forward scattering was collected at 350V and SYTO9 intensities were collected at 450V using a 488 nm excitation and 500-560 nm emission filter. For each sample, 100,000 events were recorded. Analysis was performed using the FloJo software.

### Genome sequencing and assembly

*A. B.* sp. USDA3516 was grown at 29°C overnight in a modified arabinose-gluconate medium (Sachs *et al*., 2009) with shaking. DNA was extracted from *B.* sp. USDA3516 and prepared for Oxford Nanopore sequencing, following protocols previously described, with the exception that the Ligation Sequencing Kit V14 was used (Weisberg, AJ *et al*., 2022). The *PathogenSurveillance* pipeline was used to automatically process and assemble reads, assess assembly quality, and annotate the genome sequence (Foster *et al*., 2025).

### Manual genome analysis

Preliminary annotation of the *B.* sp. USDA3516 genome was performed using Prokka version 1.14.5 (https://github.com/tseemann/prokka) (Seemann, 2014) and Bakta version 1.11.4 (https://github.com/oschwengers/bakta) (Schwengers *et al*., 2021). Protein-coding genes were annotated using the InterProScan command line tool (version 5.69-101.0, https://github.com/ebi-pf-team/interproscan) (Jones *et al*., 2014). Genes homologous with selected reference genomes were identified through custom reciprocal best BLAST hit analysis scripts using *blastp* (https://www.ncbi.nlm.nih.gov/books/NBK279690/) with an e-value threshold of 1e-20. Reference genomes selected for this analysis were *Bradyrhizobium* sp. ORS 278 (https://www.uniprot.org/taxonomy/114615), *Bradyrhizobium* BTAi1 (https://www.uniprot.org/taxonomy/288000), *Bradyrhizobium diazoefficiens* USDA110 (https://www.uniprot.org/taxonomy/224911), *Rhodopseudomonas palustris* BAA-98 (https://www.uniprot.org/taxonomy/258594), *Sinorhizobium meliloti* 1021 (https://www.uniprot.org/taxonomy/266834), *Caulobacter vibroides* (formerly *crescentus*) CB15N (https://www.uniprot.org/taxonomy/565050), *Bacillus subtilis* 168 (https://www.uniprot.org/taxonomy/224308), *Escherichia coli* K12 (https://www.uniprot.org/taxonomy/83333). Final gene annotations were made by majority-rule across the above preliminary annotation sources and reviewed manually. Chromosome and operon annotations were visualized using Proksee (https://proksee.ca/) (Grant *et al*., 2023).

Orthologous protein groups were identified using Proteinortho version 6.3.6 (https://gitlab.com/paulklemm_PHD/proteinortho) (Klemm *et al*., 2023). The following proteome accessions were used for comparison with *B.* sp USDA3516: *Bradyrhizobium arachidis* CCBAU 051107 (UniProt UP000594015), *Bradyrhizobium brasilense* R5 (UniProt UP001221546), *Bradyrhizobium* BTAi1 (UniProt UP000000246), *Bradyrhizobium cajani* 1010 (UniProt UP000449969), *Bradyrhizobium canariense* BTA-1 (UniProt UP000887172), *Bradyrhizobium diazoefficiens* USDA110 (UniProt UP000002526), *Bradyrhizobium* sp. DOA9 (NCBI RefSeq GCF_000617845.2), *Bradyrhizobium elkanii* USDA 76 (NCBI Assembly GCA_023278185.1), *Bradyrhizobium japonicum* USDA 6 (UniProt UP000005663), *Bradyrhizobium* sp. ORS 285 (UniProt UP000196578), *Bradyrhizobium semiaridum* WSM 1704 (NCBI Assembly GCA_020329505.1), *Bradyrhizobium stylosanthis* (UniProt UP000319949).

### Phylogenetic tree reconstruction

For the *Bradyrhizobium* species tree, a bac120 alignment of *Bradyrhizobium* species representatives from the Genome Taxonomy Database (GTDB) was generated using GTDB Toolkit v2.4.0+ (gtdb-tk, https://github.com/Ecogenomics/GTDBTk) (Chaumeil *et al*., 2022). For protein trees, sequences were aligned using the MUSCLE v. 5.1 sequence alignment program (https://www.drive5.com/muscle/muscle.html) (Edgar, 2004). Maximum likelihood trees were generated using IQ-TREE 2 (https://iqtree.github.io/) (Minh *et al*., 2020). The optimal nucleotide substitution model for each alignment was determined by ModelFinder (Kalyaanamoorthy *et al*., 2017) and 1,000 replicates were used to calculate branch support values using ultrafast bootstrap approximation (UFBoot) (Hoang *et al*., 2017). Trees were visualized and annotated using TreeViewer (https://treeviewer.org/) (Bianchini & Sánchez-Baracaldo, 2024).

## DISCUSSION

*A. americana* is a basal, diploid member of the Dalbergioids with potential use as a genetic model for Nod factor-dependent Dalbergioid symbiosis. Here we evaluated the symbiont range of *A. americana* from central Florida and characterized a nodule isolate from the same region, *B.* sp. USDA3516, which appears to be a close relative of the effective Thai *A. americana* symbiont *B.* sp DOA9. We find that *A. americana* is effectively nodulated by diverse non-native *Bradyrhizobium* strains but is not compatible with non-photosynthetic strains, consistent with earlier work. Microscopy of nodule cross-sections and isolated bacteroids revealed that most of the compatible symbionts of *A. americana* exhibit hallmarks of terminal bacteroid differentiation (branched morphologies and higher ploidy than cultured cells), albeit to varying degrees across symbiont species.

What is the role of morphological changes in bacteroids? Prior studies by Lamouche *et al*. (Lamouche *et al*., 2019a; Lamouche *et al*., 2019b) attempted to address this question by inoculating the same set of *Bradyrhizobium* strains (*Bradyrhizobium* sp. ORS285, ORS287, ORS335, and ORS357) onto three jointvetch hosts: two species that induce spherical bacteroids (*A. evenia* and *A. indica*) and one species with elongated bacteroids (*A. afraspera*). The authors then quantified the benefit conferred to each plant by each symbiont via the increase in plant biomass relative to the nodule biomass – a proxy for the benefit to the host relative to the host’s investment in symbiosis. Using this metric, the two jointvetches inducing spherical bacteroids were consistently observed to receive greater benefits from the same strains than the jointvetch with elongated bacteroids. Based on this, the authors concluded that spherical bacteroids may be inherently more effective than elongated bacteroids.

Our study contradicts the view that bacteroid morphology *per se* is a major determinant of symbiotic efficiency. Within the same host, we identified symbionts with four distinct bacteroid phenotypes: (i) branching only, (ii) elevated ploidy only, (iii) both branching and elevated ploidy, and (iv) neither branching nor elevated ploidy. We found that branching alone was not sufficient for a strain to function as an effective symbiont, whereas elevated ploidy was observed in all effective strains. Though the low number of effective symbionts that we identified does not provide a large sample to suggest that higher bacteroid ploidy is strictly necessary for effective *A. americana* symbiosis, we can conclude that branched bacteroid morphologies are not required, a conclusion that is also supported by the effective interaction between *A. americana* and the non-branching bacteroids of *B.* sp. DOA9 (Noisangiam *et al*., 2012).

The disagreement between this work and Lamouche *et al*. has several potential explanations. Our groups employ different metrics for evaluating symbiosis (proportion of non-nodule biomass conferred vs. acetylene reduction), and the significance of bacteroid morphological changes may differ between *A. americana* and the other jointvetches examined. However, we note that Lamouche *et al*. rely on comparison across rather than within jointvetch species, and that species-specific differences in symbiotic mechanisms beyond morphological changes may account for the symbiosis efficiencies in spherical bacteroid- vs. elongated bacteroid-producing hosts. Ultimately, a careful examination of bacteroid morphotype relevance will require generating symbiont strains lacking only the factors required for bacteroid branching, which could then be assayed in a single host-symbiont context.

Currently we do not know what genetic factors regulate morphological and/or ploidy changes in *A. americana* bacteroids, nor in those of most other Dalbergioid legumes. *A. americana* presumably produces NCR peptides, and the presence of branched bacteroids likely relates to the degree of compatibility between *A. americana* NCRs and the presence or sequence of a symbiont’s cell cycle control factors. Others have found that disruption of the cell division factor FtsZ in species of Hyphomicrobiales/Rhizobiales order is sufficient to induce branched cells in culture (Aubry *et al*., 2025), as is overexpression of Min proteins *Sinorhizobium meliloti* (where Min proteins are non-essential) (Cheng *et al*., 2007). It is curious, then, that we identified a *minCDE-*adjacent operon that is conserved in all *A. americana* symbionts with branched bacteroids, though the function of these genes – and whether they affect Min protein levels – is unknown. This provides the foundation for future work to determine whether *A. americana* indeed produces NCR peptides, what these peptides target, and whether the conserved *minCDE* genomic region is relevant to bacteroid differentiation.

## ACKNOWLEDGEMENTS

Funding for this project was provided by the Carnegie Institution for Science Endowment to B.J.B. Pilot experiments were performed by students in the Marine Biological Laboratory’s Molecular & Cell Biology of Symbiosis course, which is funded by a grant from the Gordon & Betty Moore Foundation. We thank Will Ludington (Johns Hopkins University) for Attune flow cytometry access and members of the Ludington lab for training and technical assistance. Members of the Belin lab provided helpful discussions of the manuscript, and Mahmud Siddiqi (Carnegie) provided technical assistance for microscopy. We are grateful to Carnegie Embryology’s IT, front office, and facilities support staff for making our work possible.

## AUTHOR CONTRIBUTIONS

T.S.C., A.A.A., and A.S. performed *A. americana* inoculation experiments and bacteroid analyses. J.R.S. sequenced and assembled the *B.* sp. USDA3516 genome. C.P.R., R.A.B., J.H.C., and B.J.B. analyzed the *B.* sp. USDA3516 genome. B.J.B. designed the project, acquired funding, and prepared the manuscript.

## DATA AVAILABIITY

*Bradyrhizobium* sp. USDA3615 genome assembly information is available on NCBI, accession #**XXXXXXX**.

---

**Figure S1.**
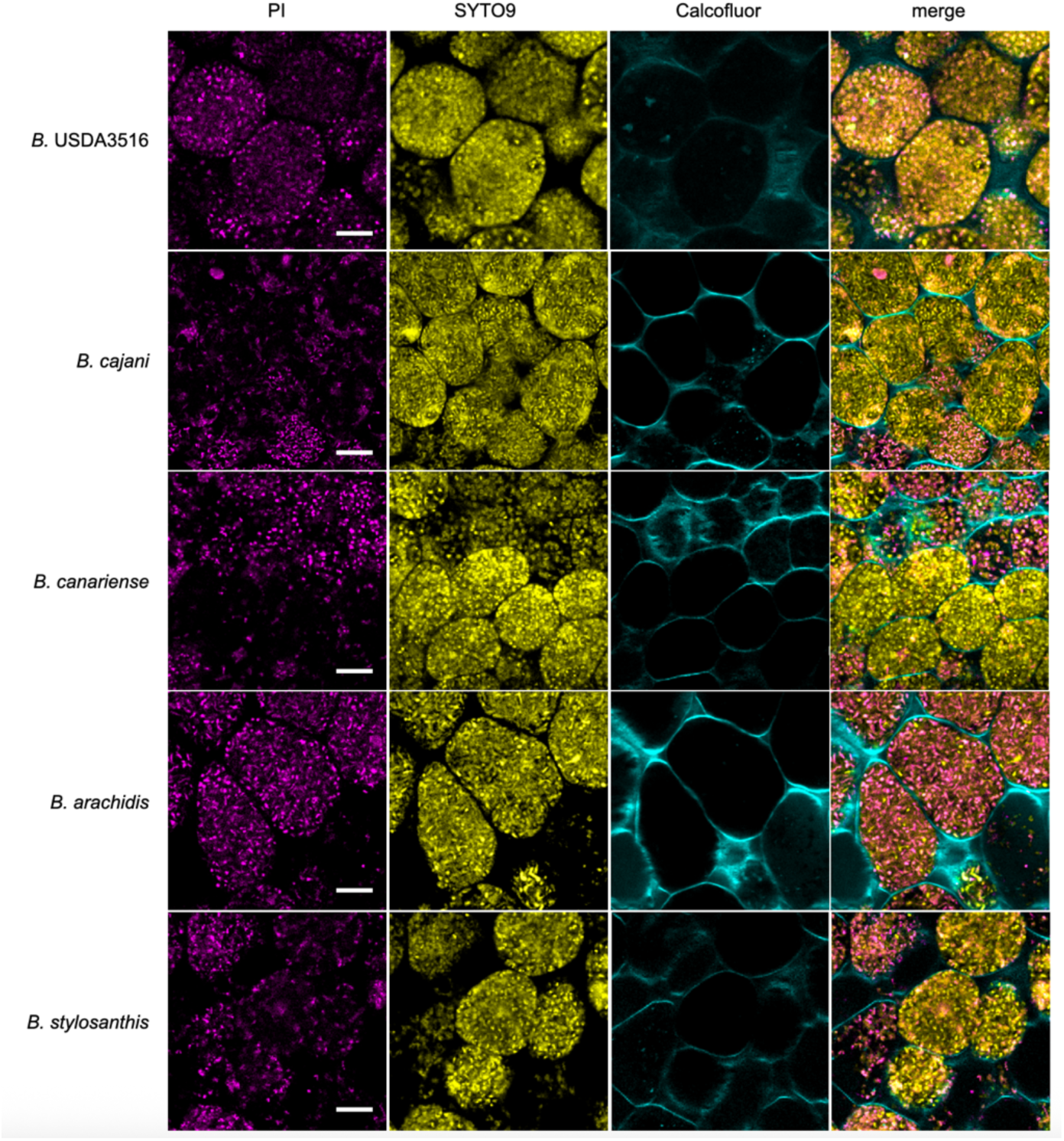
Representative confocal images of nodule cross-sections collected at 35 dpi. Sections were stained with Calcofluor (cyan), SYTO9 (yellow), and propidium iodide (“PI”; magenta). Scale bars = 10 microns.

**Figure S2.**
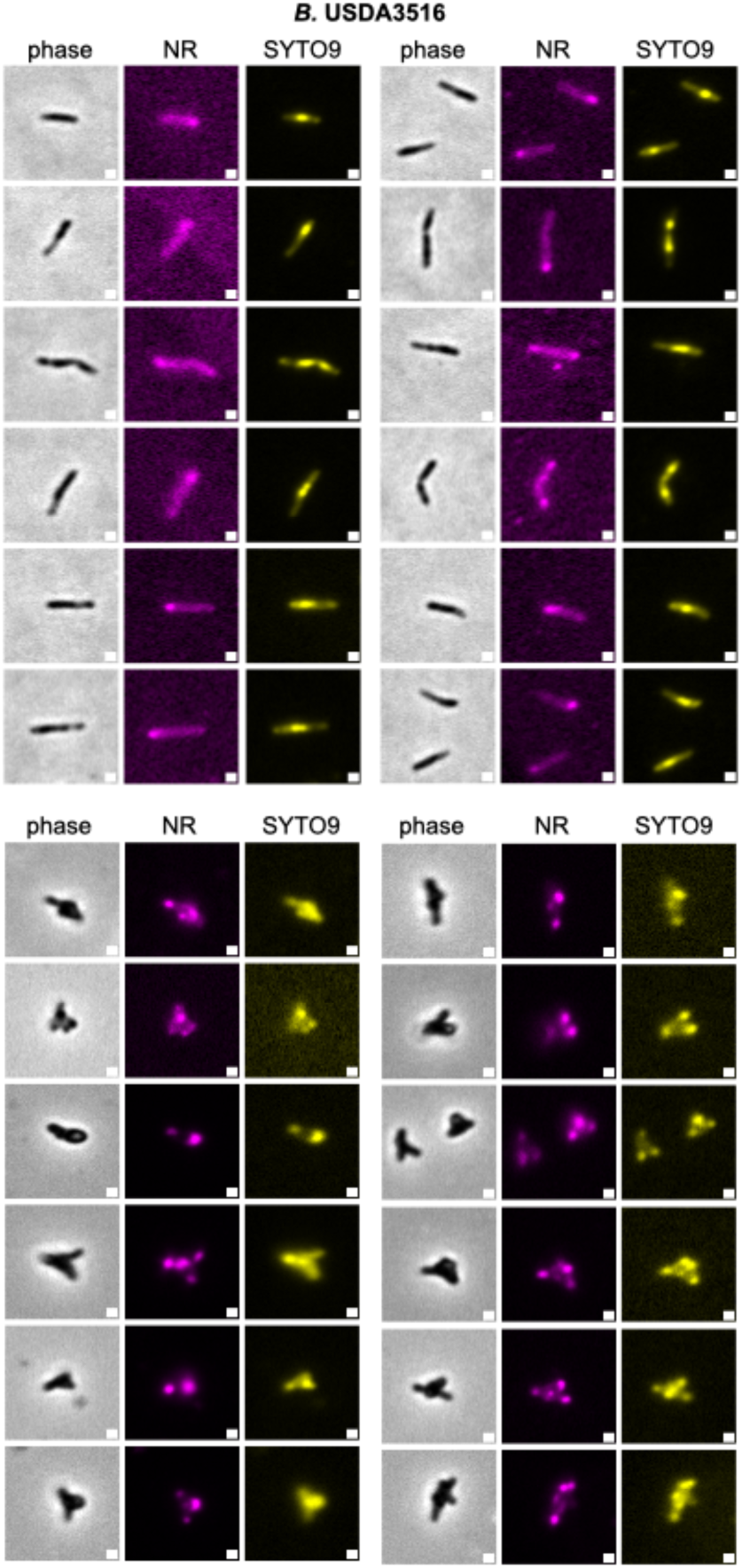
Additional phase and fluorescence images for *B.* sp. USDA3516 bacteroids and cultured cells stained with Nile Red (NR) and SYTO9. Scale bars indicate 1 micron.

**Figure S3.**
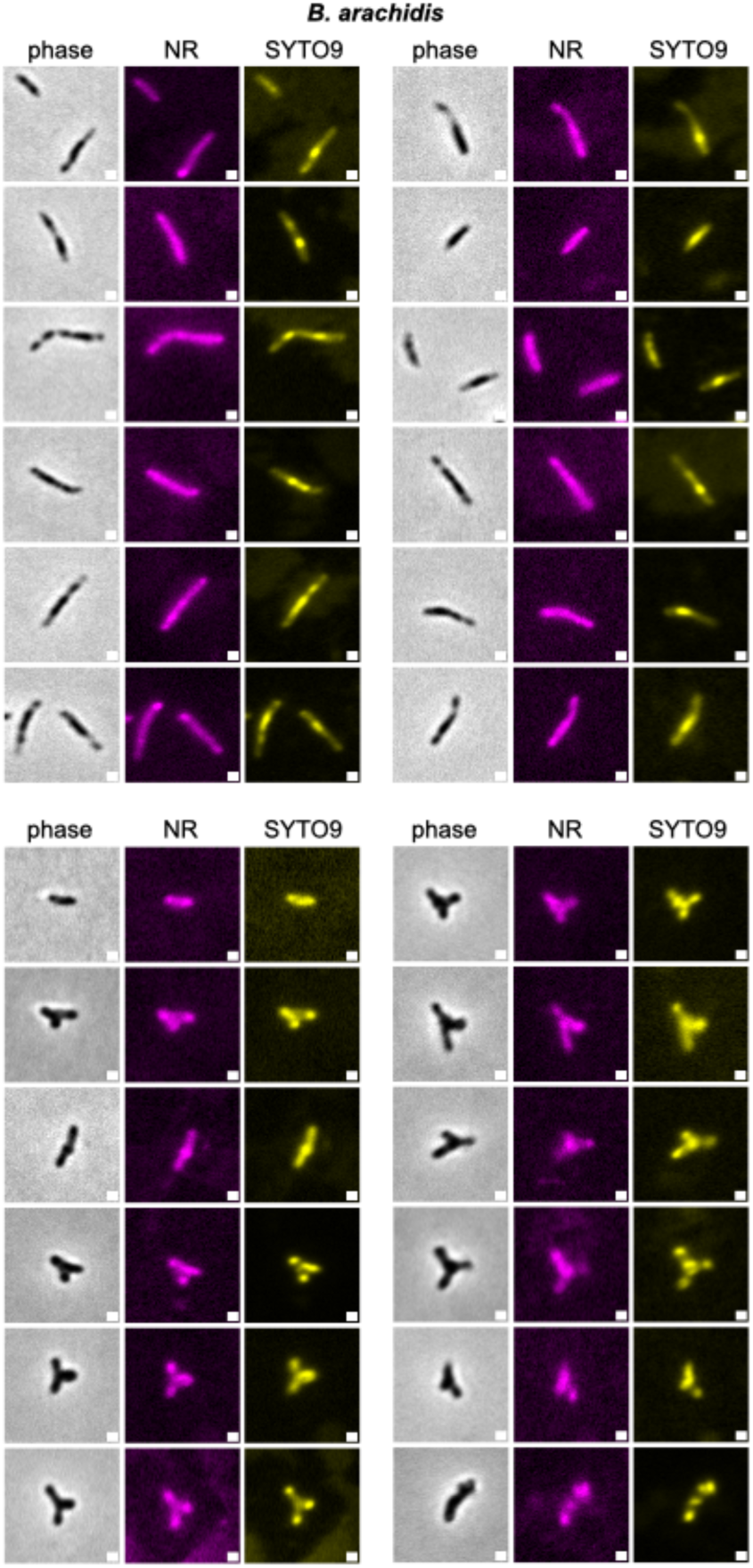
Additional phase and fluorescence images for *B. arachidis* bacteroids and cultured cells stained with Nile Red (NR) and SYTO9. Scale bars indicate 1 micron.

**Figure S4.**
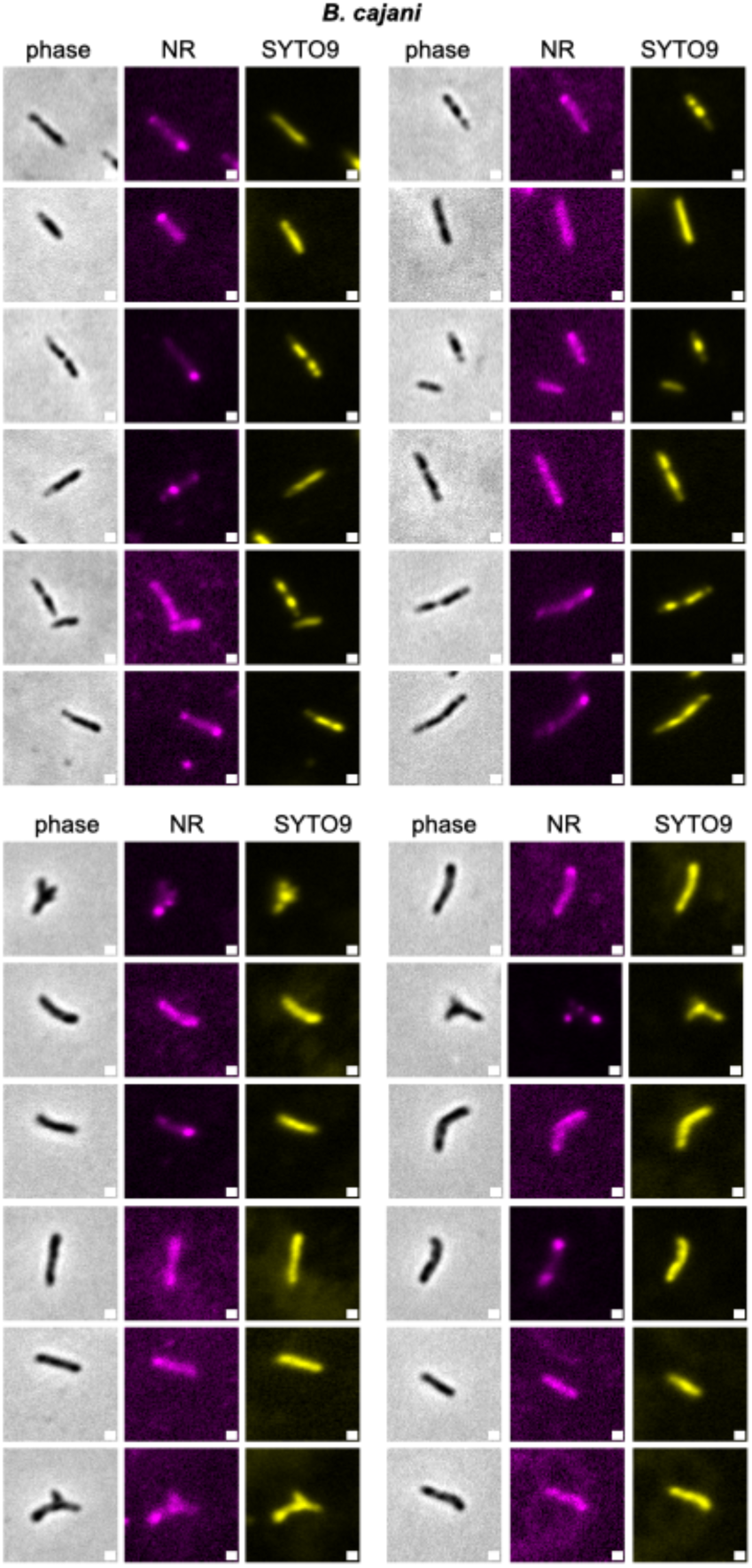
Additional phase and fluorescence images for *B. cajani* bacteroids and cultured cells stained with Nile Red (NR) and SYTO9. Scale bars indicate 1 micron.

**Figure S5.**
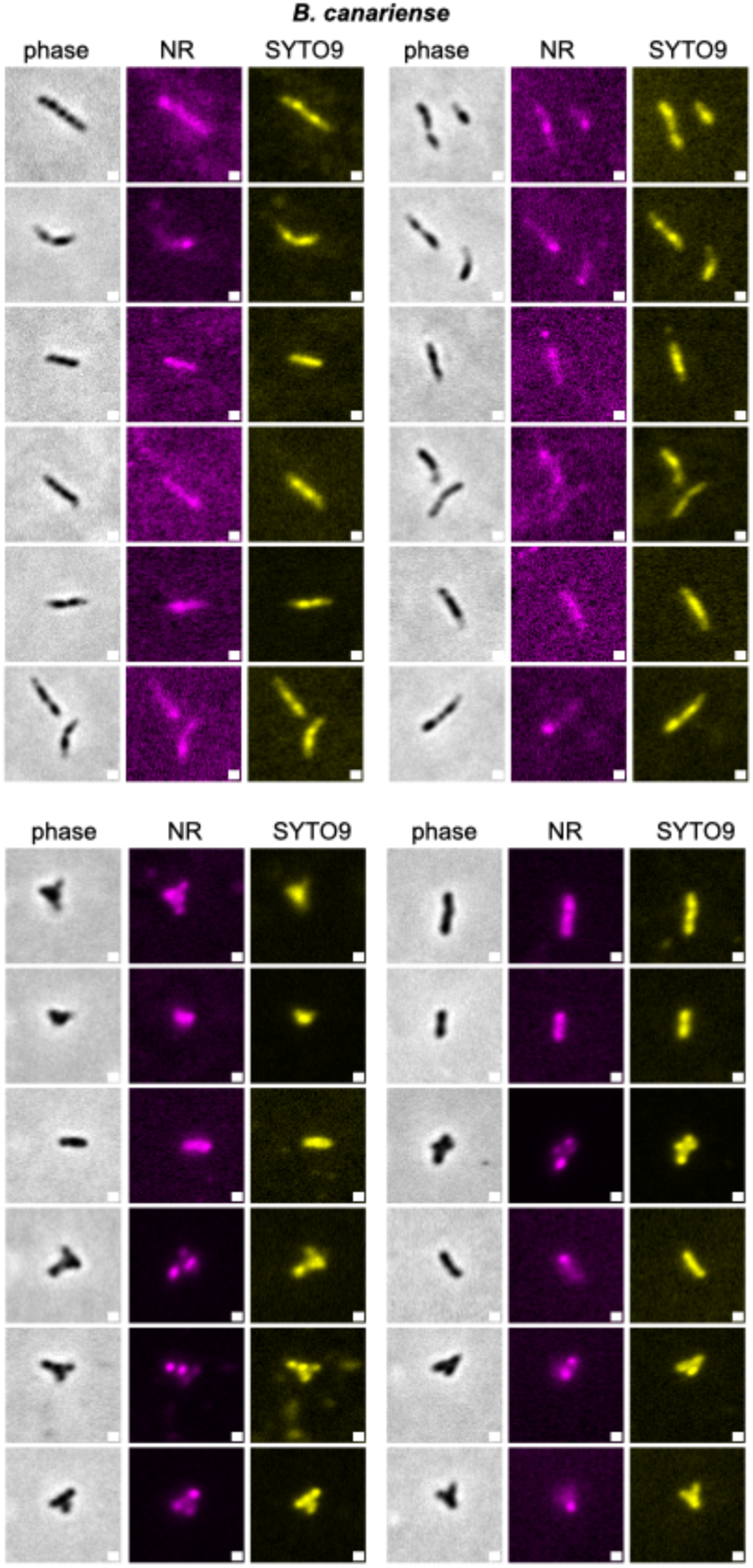
Additional phase and fluorescence images for *B. canariense* bacteroids and cultured cells stained with Nile Red (NR) and SYTO9. Scale bars indicate 1 micron.

**Figure S6.**
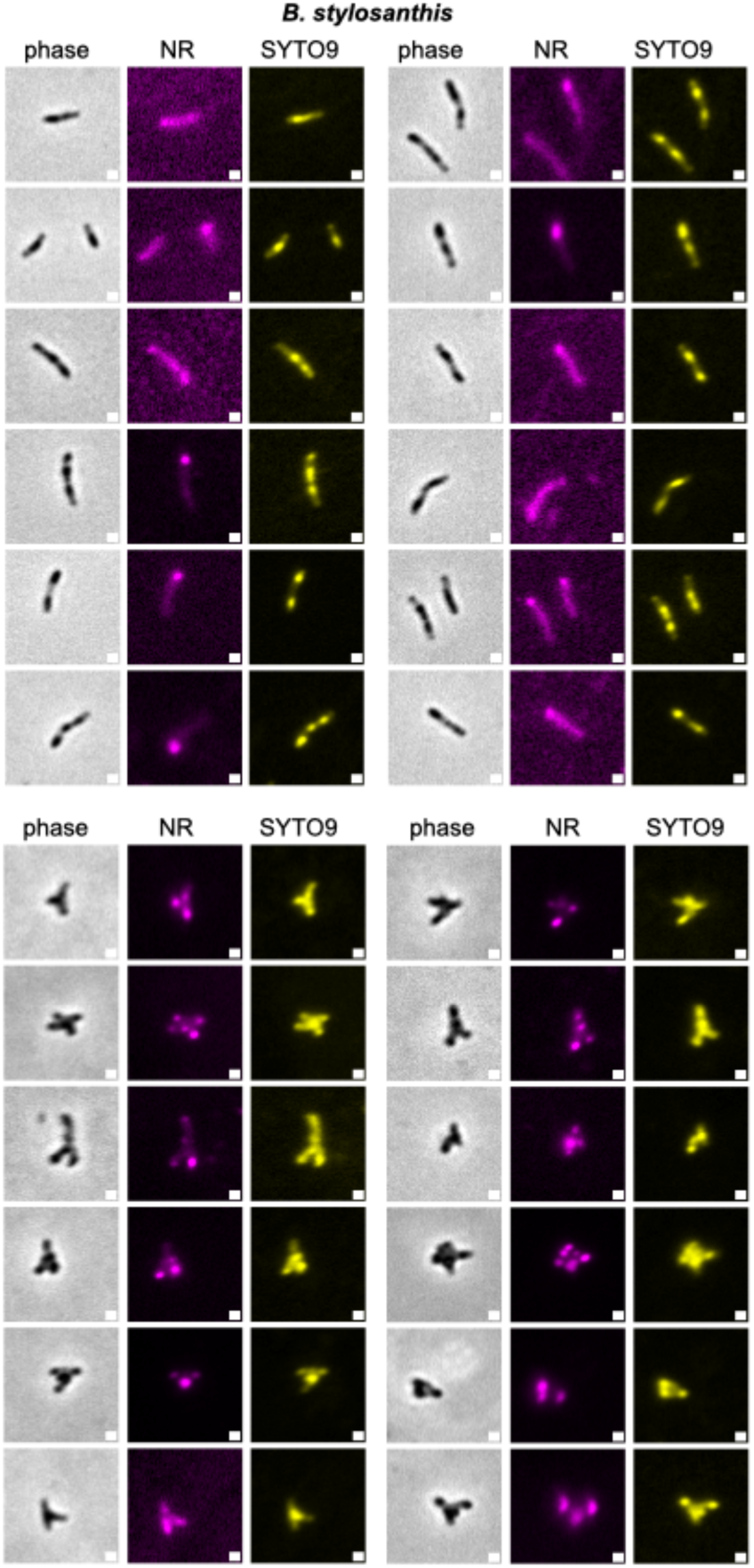
Additional phase and fluorescence images for *B. stylosanthis* bacteroids and cultured cells stained with Nile Red (NR) and SYTO9. Scale bars indicate 1 micron.

**Figure S7.**
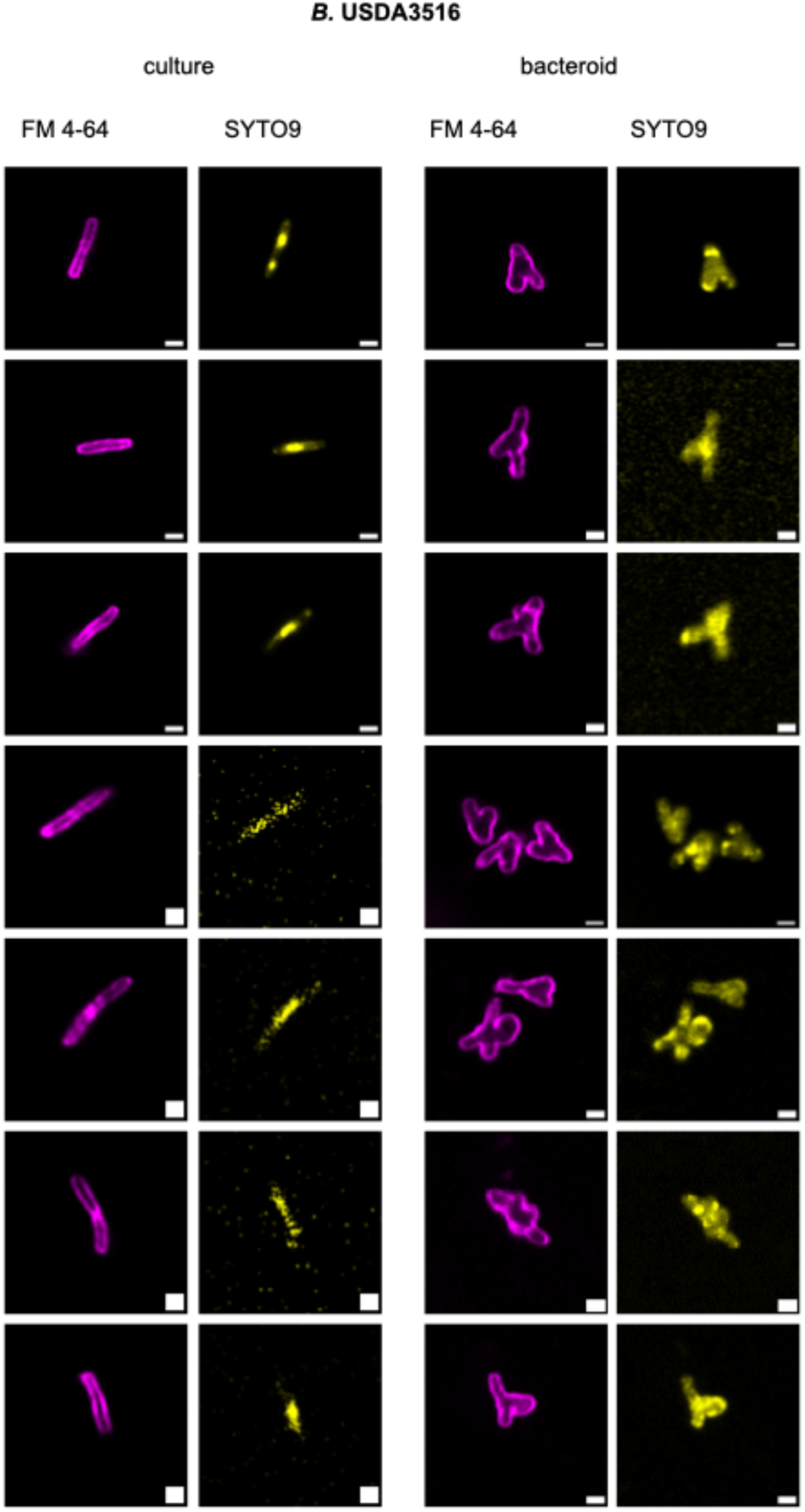
Airyscan superresolution imaging of *B.* sp. USDA3516 bacteroids and cultured cells stained with FM 4- 64 and SYTO9. Scale bars indicate 1 micron.

**Figure S8.**
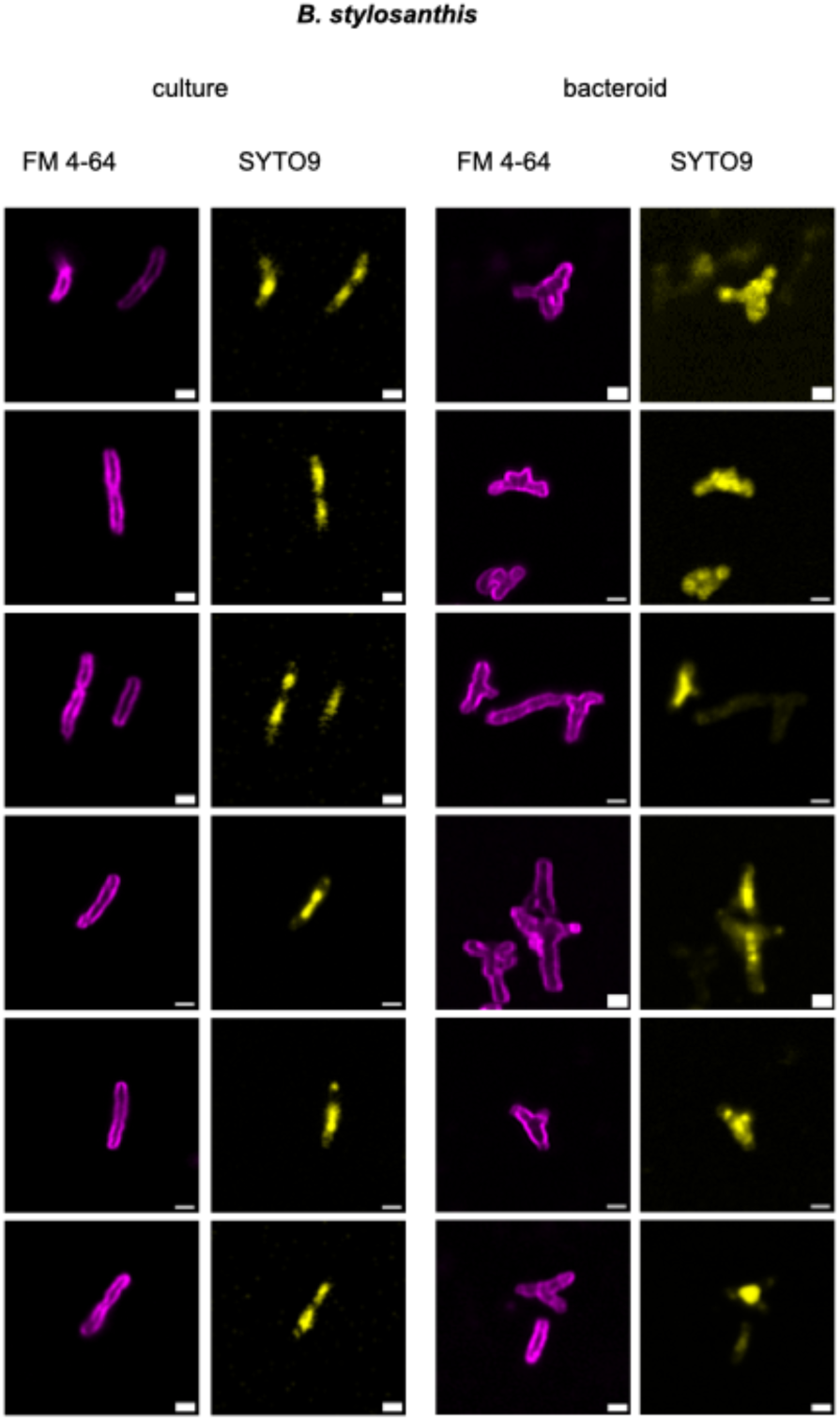
Airyscan superresolution imaging of *B. stylosanthis* bacteroids and cultured cells stained with FM 4-64 and SYTO9. Scale bars indicate 1 micron.

**Figure S9.**
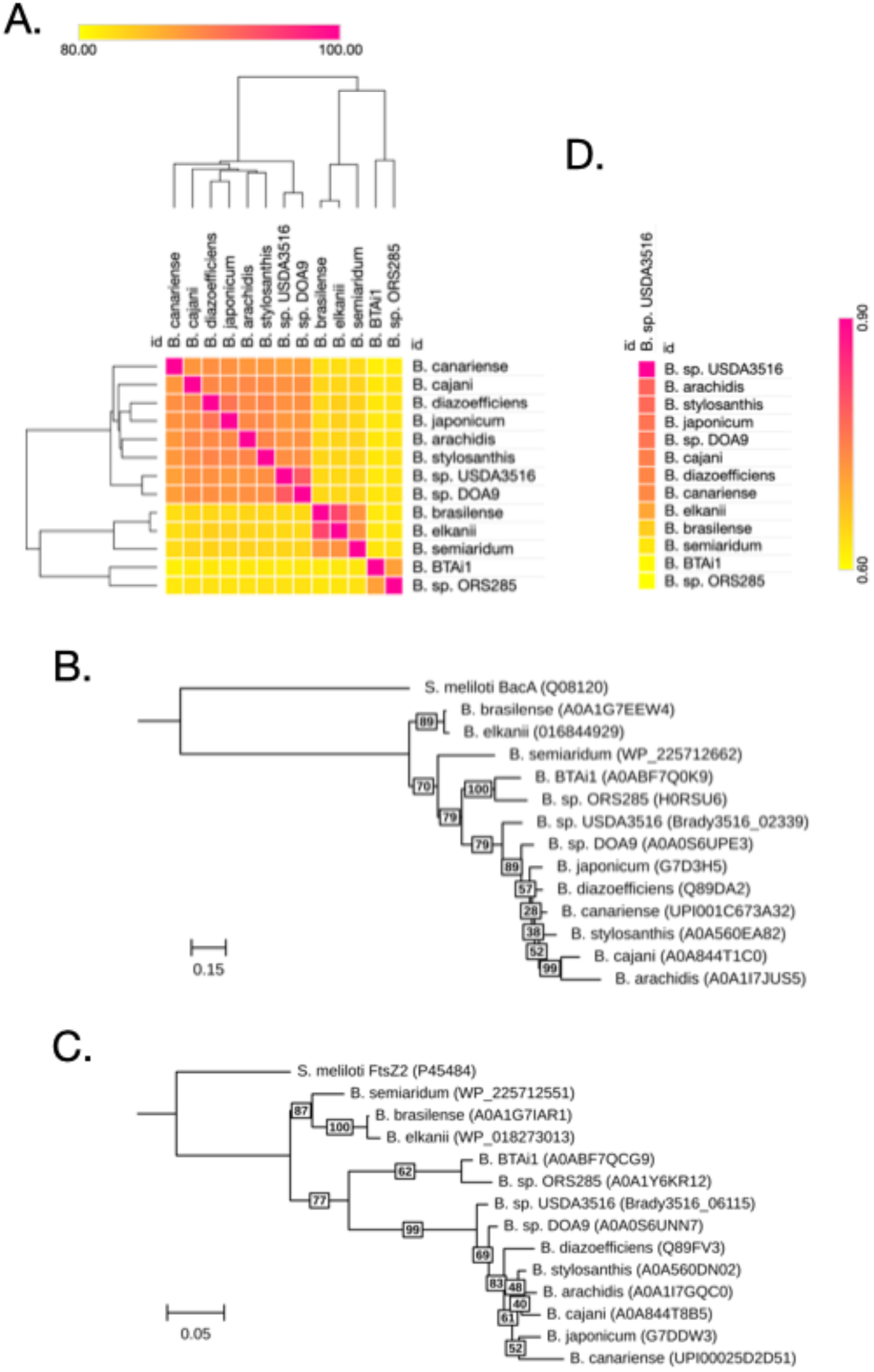
(A) Heatmap of average nucleotide identity across stains used for *A. americana* inoculation and *B.* sp. DOA9. Calculated with FastANI v. 1.34 (https://github.com/ParBLiSS/FastANI) (Jain *et al*., 2018) and visualized with Morpheus (https://software.broadinstitute.org/morpheus/) (B-C) Maximum likelihood phylogenetic tree of BclA (B) and FtsZ (C) homologs from genomes in (A). Numbers at internal nodes (highlighted in white) indicate branch support values using ultrafast bootstrap approximation (UFBoot), based on 1000 bootstraps: 100 = highest confidence, 0 = no confidence. UniProt IDs are provided in parentheses in the branch labels. Outgroup sequences are *S. meliloti* homologs. Scale bars indicate substitutions per 1000 amino acids. (D) Heatmap illustrating the percentage of orthologous gene groups (OGs) shared between *B.* sp. 3516 and all other strains in (A).

**Table S1.** Manually curated gene annotations in *B.* sp. USDA3516. Information provided in each column follows: (A) Ordered locus ID, (B) Contig, (C-D) Start and stop nucleotide positions, relative to the contig; (E) Coding strand (+ or -); (F) Preferred gene names, where available; (G) Consensus gene description; (H) Prokka suggested gene name; (I) Prokka suggested gene description; (J) Bakta suggested gene name; (K) Bakta suggested gene description; (L) InterPro domains identified by Interproscan, formatted as Domain ID:Domain Name; (M) Gene name of reciprocal best BLAST hit (RBBH) from *Bradyrhizobium* sp. ORS 278, when present; (N) Gene description of RBBH from *Bradyrhizobium* sp. ORS 278, when present; (O) Gene name of RBBH from *Bradyrhizobium* BTAi1, when present; (P) Gene description of RBBH from *Bradyrhizobium* BTAi1, when present; (Q) Gene name of RBBH from *Bradyrhizobium diazoefficiens* USDA110, when present; (R) Gene description of RBBH from *Bradyrhizobium diazoefficiens* USDA110, when present; (S) Gene name of RBBH from *Rhodopseudomonas palustris* BAA-98, when present; (T) Gene description of RBBH from *Rhodopseudomonas palustris* BAA-98, when present; (U) Gene name of RBBH from *Sinorhizobium meliloti* 1021, when present; (V) Gene description of RBBH from *Sinorhizobium meliloti* 1021, when present; (W) Gene name of RBBH from *Caulobacter vibroides* (formerly *crescentus*) CB15N, when present; (X) Gene description of RBBH from *Caulobacter vibroides* (formerly *crescentus*) CB15N, when present; (Y) Gene name of RBBH from *Bacillus subtilis* 168, when present; (Z) Gene description of RBBH from *Bacillus subtilis* 168, when present; (AA) Gene name of RBBH from *Escherichia coli* K12, when present; (BB) Gene description of RBBH from *Escherichia coli* K12, when present.

## REFERENCES

1. Aubry B, Randich A, Hudson B, Horton E, Brown PJB. 2025. Ancient Truncated FtsZ Paralogs Likely Tune Cell Division in Hyphomicrobiales. bioRxiv: 2025.2009.2029.679267.

2. Bianchini G, Sánchez-Baracaldo P. 2024. TreeViewer: Flexible, modular software to visualise and manipulate phylogenetic trees. Ecology and Evolution 14(2).

3. Bonaldi K, Gargani D, Prin Y, Fardoux J, Gully D, Nouwen N, Goormachtig S, Giraud E. 2011. Nodulation of Aeschynomene afraspera and A. indica by Photosynthetic Bradyrhizobium Sp. Strain ORS285: The Nod-Dependent Versus the Nod-Independent Symbiotic Interaction. Molecular Plant-Microbe Interactions 24(11): 1359–1371.

4. Boukherissa A, Sankari S, Timchenko T, Bourge M, Mergaert P, diCenzo GC, Shykoff JA, Alunni B, Rodríguez de la Vega RC. 2025. Structure-based phylogenetic analysis reveals multiple events of convergent evolution of cysteine-rich antimicrobial peptides in legume-rhizobium symbiosis. bioRxiv: 2025.2009.2009.675119.

5. Brottier L, Chaintreuil C, Simion P, Scornavacca C, Rivallan R, Mournet P, Moulin L, Lewis G, Fardoux J, Brown S, et al. 2018. A phylogenetic framework of the legume genus Aeschynomene for comparative genetic analysis of the Nod-dependent and Nod-independent symbioses. Bmc Plant Biology 18.

6. Chaintreuil C, Gully D, Hervouet C, Tittabutr P, Randriambanona H, Brown S, Lewis G, Bourge M, Cartieaux F, Boursot M, et al. 2016. The evolutionary dynamics of ancient and recent polyploidy in the African semiaquatic species of the legume genus Aeschynomene. New Phytologist 211(3): 1077– 1091.

7. Chaumeil P-A, Mussig AJ, Hugenholtz P, Parks DH. 2022. GTDB-Tk v2: memory friendly classification with the genome taxonomy database. Bioinformatics 38(23): 5315–5316.

8. Cheng J, Sibley C, Zaheer R, Finan T. 2007. A Sinorhizobium meliloti minE mutant has an altered morphology and exhibits defects in legume symbiosis. Microbiology-Sgm 153: 375–387.

9. Czernic P, Gully D, Cartieaux F, Moulin L, Guefrachi I, Patrel D, Pierre O, Fardoux J, Chaintreuil C, Nguyen P, et al. 2015. Convergent Evolution of Endosymbiont Differentiation in Dalbergioid and Inverted Repeat-Lacking Clade Legumes Mediated by Nodule-Specific Cysteine-Rich Peptides. Plant Physiology 169(2): 1254–1265.

10. Edgar RC. 2004. MUSCLE: multiple sequence alignment with high accuracy and high throughput. Nucleic Acids Research 32(5): 1792–1797.

11. Foster ZSL, Sudermann MA, Parada Rojas CH, Blair LK, Iruegas Bocardo F, Dhakal U, Weisberg AJ, Phan H, Chang JH, Grunwald NJ. 2025. PathogenSurveillance: an automated pipeline for population genomic analyses and pathogen identification. bioRxiv: 2025.2010.2031.685798.

12. Garrido-Sanz D, Redondo-Nieto M, Mongiardini E, Blanco-Romero E, Durán D, Quelas J, Martin M, Rivilla R, Lodeiro A, Althabegoiti M. 2019. Phylogenomic Analyses of Bradyrhizobium Reveal Uneven Distribution of the Lateral and Subpolar Flagellar Systems, Which Extends to Rhizobiales. Microorganisms 7(2).

13. Giraud E, Moulin L, Vallenet D, Barbe V, Cytryn E, Avarre J, Jaubert M, Simon D, Cartieaux F, Prin Y, et al. 2007. Legumes symbioses:: Absence of Nod genes in photosynthetic bradyrhizobia. Science 316(5829): 1307–1312.

14. Grant J, Enns E, Marinier E, Mandal A, Herman E, Chen C, Graham M, Van Domselaar G, Stothard P. 2023. Proksee: in-depth characterization and visualization of bacterial genomes. Nucleic Acids Research 51(W1): W484–W492.

15. Grant W, Trese A. 1996. Developmental regulation of nodulation in Arachis hypogea (peanut) and Aeschynomene americana (jointvetch). Symbiosis 20(3): 247–258.

16. Guefrachi I, Pierre O, Timchenko T, Alunni B, Barrière Q, Czernic P, Villaécija-Aguilar J, Verly C, Bourge M, Fardoux J, et al. 2015. Bradyrhizobium BclA Is a Peptide Transporter Required for Bacterial Differentiation in Symbiosis with Aeschynomene Legumes. Molecular Plant-Microbe Interactions 28(11): 1155–1166.

17. Guerra-Garcia F, Sankari S. 2025. NCR peptides in plant-bacterial symbiosis: applications and importance. Trends in Microbiology 33(2): 147–150.

18. Guha S, Molla F, Sarkar M, Ibañez F, Fabra A, DasGupta M. 2022. Nod factor-independent ’crack-entry’ symbiosis in dalbergoid legume Arachis hypogaea. Environmental Microbiology 24(6): 2732–2746.

19. Gully D, Czernic P, Cruveiller S, Mahé F, Longin C, Vallenet D, François P, Nidelet S, Rialle S, Giraud E, et al. 2018. Transcriptome Profiles of Nod Factor-independent Symbiosis in the Tropical Legume Aeschynomene evenia. Scientific Reports 8.

20. Hoang DT, Chernomor O, von Haeseler A, Minh BQ, Vinh LS. 2017. UFBoot2: Improving the Ultrafast Bootstrap Approximation. Molecular Biology and Evolution 35(2): 518–522.

21. Jain C, Rodriguez-R LM, Phillippy AM, Konstantinidis KT, Aluru S. 2018. High throughput ANI analysis of 90K prokaryotic genomes reveals clear species boundaries. Nature Communications 9(1): 5114.

22. Jones P, Binns D, Chang H, Fraser M, Li W, McAnulla C, McWilliam H, Maslen J, Mitchell A, Nuka G, et al. 2014. InterProScan 5: genome-scale protein function classification. Bioinformatics 30(9): 1236–1240.

23. Kalyaanamoorthy S, Minh BQ, Wong TKF, von Haeseler A, Jermiin LS. 2017. ModelFinder: fast model selection for accurate phylogenetic estimates. Nature Methods 14(6): 587–589.

24. Klemm P, Stadler P, Lechner M. 2023. Proteinortho6: pseudo-reciprocal best alignment heuristic for graph-based detection of (co-)orthologs. Frontiers in Bioinformatics 3.

25. Lamouche F, Bonadé-Bottino N, Mergaert P, Alunni B. 2019a. Symbiotic Efficiency of Spherical and Elongated Bacteroids in the Aeschynomene-Bradyrhizobium Symbiosis. Frontiers in Plant Science 10.

26. Lamouche F, Gully D, Chaumeret A, Nouwen N, Verly C, Pierre O, Sciallano C, Fardoux J, Jeudy C, Szucs A, et al. 2019b. Transcriptomic dissection of Bradyrhizobium sp. strain ORS285 in symbiosis with Aeschynomene spp. inducing different bacteroid morphotypes with contrasted symbiotic efficiency. Environmental Microbiology 21(9): 3244–3258.

27. Legume Phylogeny Working Group (LPWG) A, G. C., Atahuachi Burgos, M., Bagnatori Sartori, Â. L., Balan, A., Bandyopadhyay, S., Barbosa Pinto, R., Barrett, R., Boatwright, J. S., Borges, L. M., Bortoluzzi, R., Broich, S. L., Brullo, S., Bruneau, A., Cardinal-McTeague, W., Cardoso, D., Castro Silva, I. C., Cervantes, A., Choo, L. M., et al. . 2025. The World Checklist of Vascular Plants (WCVP): Fabaceae: Royal Botanic Gardens, Kew, Richmond, UK.

28. Minh BQ, Schmidt HA, Chernomor O, Schrempf D, Woodhams MD, von Haeseler A, Lanfear R. 2020. IQ-TREE 2: New Models and Efficient Methods for Phylogenetic Inference in the Genomic Era. Molecular Biology and Evolution 37(5): 1530–1534.

29. Mornico D, Miché L, Béna G, Nouwen N, Verméglio A, Vallenet D, Smith A, Giraud E, Médigue C, Moulin L. 2012. Comparative Genomics of Aeschynomene Symbionts: Insights into the Ecological Lifestyle of Nod-Independent Photosynthetic Bradyrhizobia. Genes 3(1): 35–61.

30. Nations FAOotU. 2024. Agricultural production statistics 2010–2023. FAOSTAT Analytical Briefs. Rome, Italy.

31. Noisangiam R, Teamtisong K, Tittabutr P, Boonkerd N, Toshiki U, Minamisawa K, Teaumroong N. 2012. Genetic Diversity, Symbiotic Evolution, and Proposed Infection Process of Bradyrhizobium Strains Isolated from Root Nodules of Aeschynomene americana L. in Thailand. Applied and Environmental Microbiology 78(17): 6236–6250.

32. Okazaki S, Noisangiam R, Okubo T, Kaneko T, Oshima K, Hattori M, Teamtisong K, Songwattana P, Tittabutr P, Boonkerd N, et al. 2015. Genome Analysis of a Novel Bradyrhizobium sp DOA9 Carrying a Symbiotic Plasmid. Plos One 10(2).

33. Pan H, Shim A, Lubin M, Belin B. 2024. Hopanoid lipids promote soybean-Bradyrhizobium symbiosis. Mbio 15(4).

34. Pessi G, Ahrens C, Rehrauer H, Lindemann A, Hauser F, Fischer H, Hennecke H. 2007. Genome-wide transcript analysis of Bradyrhizobium japonicum bacteroids in soybean root nodules. Molecular Plant-Microbe Interactions 20(11): 1353–1363.

35. Quilbé J, Lamy L, Brottier L, Leleux P, Fardoux J, Rivallan R, Benichou T, Guyonnet R, Becana M, Villar I, et al. 2021. Genetics of nodulation in Aeschynomene evenia uncovers mechanisms of the rhizobium-legume symbiosis. Nature Communications 12(1).

36. Raul B, Bhattacharjee O, Ghosh A, Upadhyay P, Tembhare K, Singh A, Shaheen T, Ghosh A, Torres-Jerez I, Krom N, et al. 2022. Microscopic and Transcriptomic Analyses of Dalbergoid Legume Peanut Reveal a Divergent Evolution Leading to Nod-Factor-Dependent Epidermal Crack-Entry and Terminal Bacteroid Differentiation. Molecular Plant-Microbe Interactions 35(2): 131–145.

37. Sachs JL, Kembel SW, Lau AH, Simms EL. 2009. In Situ Phylogenetic Structure and Diversity of Wild Bradyrhizobium Communities. Applied and Environmental Microbiology 75(14): 4727–4735.

38. Schwengers O, Jelonek L, Dieckmann M, Beyvers S, Blom J, Goesmann A. 2021. Bakta: rapid and standardized annotation of bacterial genomes via alignment- free sequence identification. Microbial Genomics 7(11).

39. Seemann T. 2014. Prokka: rapid prokaryotic genome annotation. Bioinformatics 30(14): 2068–2069.

40. Teamtisong K, Songwattana P, Noisangiam R, Piromyou P, Boonkerd N, Tittabutr P, Minamisawa K, Nantagij A, Okazaki S, Abe M, et al. 2014. Divergent Nod-Containing Bradyrhizobium sp DOA9 with a Megaplasmid and its Host Range. Microbes and Environments 29(4): 370–376.

41. Weisberg A, Sachs J, Chang J. 2022. Dynamic Interactions Between Mega Symbiosis ICEs and Bacterial Chromosomes Maintain Genome Architecture. Genome Biology and Evolution 14(6).

42. Weisberg AJ, Rahman A, Backus D, Tyavanagimatt P, Chang JH, Sachs JL. 2022. Pangenome Evolution Reconciles Robustness and Instability of Rhizobial Symbiosis. Mbio 13(3): e00074–00022.

43. Willems A, Doignon-Bourcier F, Coopman R, Hoste B, de Lajudie P, Gillis M. 2000. AFLP fingerprint analysis of Bradyrhizobium strains isolated from Faidherbia albida and Aeschynomene species. Systematic and Applied Microbiology 23(1): 137–147.

44. Wojciechowski M, Lavin M, Sanderson M. 2004. A phylogeny of legumes (Leguminosae) based on analyses of the plastid matK gene resolves many well-supported subclades within the family. American Journal of Botany 91(11): 1846–1862.

